# Spheroplasted cells: a game changer for DNA delivery to diatoms

**DOI:** 10.1101/2024.10.10.617634

**Authors:** E.J.L. Walker, M. Pampuch, L. Deng., Y. Li, G. Tran, T. Mock, B.J. Karas

## Abstract

Diatoms produce 20% of the world’s fixed organic carbon annually, making them vital to global carbon fixation and climate change mitigation. Their potential as cell factories for biofuels, proteins, and other high value chemicals remains underutilized due to a lack of genetic engineering tools, with DNA delivery being one of the biggest challenges. Here, we present optimized electroporation and polyethylene glycol transformation methods for delivering DNA and ribonucleoprotein complexes to *Phaeodactylum tricornutum*, a model diatom species and emerging chassis for algal biotechnology. It was possible to recover transformants with as little as 1 ng of DNA, and to transform linear or circular episomes as large as 55.6 kb. With the optimized electroporation protocol, episomes can be assembled in the algal cell *de novo* through diatom in vivo assembly (DIVA), forgoing the need for time-consuming traditional cloning steps in *Escherichia coli* and *Saccharomyces cerevisiae*. It was also possible to electroporate a Cas9 ribonucleoprotein complex in *P. tricornutum*, providing an alternative to biolistics for DNA free genome engineering. We have demonstrated that the PEG approach can be adapted to successfully transform *Thalassiosira pseudonana*, demonstrating the applicability of our methods for engineering other diatom species. These tools can be used to accelerate diatom synthetic biology projects and, therefore, the development of sustainable technologies.

## INTRODUCTION

Climate change, population expansion, and soil depletion are among several challenges that existentially threaten current agricultural practices. Due to their comparatively rapid growth and higher photosynthetic efficiency, unicellular algae have been proposed as a sustainable agricultural alternative to traditional crops since the 1950s^1^. Diatoms (Bacillariophyta) are of particular interest for this purpose as they are widely distributed across the world’s oceans^2^, where they are one of the most photosynthetically productive taxa – accounting for an estimated 20% of total fixed organic carbon annually^3^. Despite their ubiquity and productivity, the use of diatoms as cell factories has been impeded by a lack of genetic engineering tools, with DNA delivery being a major impasse.

*Phaeodactylum tricornutum* is considered to be one of the most characterized model diatom species, having a fully sequenced telomere-to-telomere reference genome^4^ in addition to molecular tools for DNA transformation^5–8^, CRISPR-based genome engineering^9^, protein expression^10^, cloning of the whole organelle genomes^11,12^, and more. These characteristics have made *P. tricornutum* an emerging chassis for diatom biotechnology^13^ and the subject of a whole-genome synthesis effort^14^. The latter, termed the Pt-syn 1.0 project, has brought together an international consortium of phycologists with the aim of replacing all the native chromosomes in the nuclear, chloroplast, and mitochondrial genomes with synthetic counterparts. To actualize this, and to discover the full potential of diatoms as cell factories, a pipeline for efficiently and rapidly testing different DNA constructs in *P. tricornutum* must be established.

Existing methods for delivering DNA to *P. tricornutum* include biolistic bombardment^8^, electroporation^5,7^, polyethylene glycol (PEG) transformation^6^, and bacterial conjugation^6^. Biolistics was the first method developed for transforming *P. tricornutum* and has been used to integrate non-replicative DNA into the nuclear genome^8,15^, with reported efficiencies of 1 to 100 transformants per 10^8^ cells. Though reasonably efficient, this method requires the use of expensive equipment (i.e., a biolistic particle delivery system) that many labs do not have access to; comparatively, conjugation, electroporation, and PEG transformation can all be conducted with more standard equipment. Of these, conjugation is the most efficient transformation method for delivering episomes (4 transformants per 10^-4^ cells)^6^, but also the most mutation prone^6,9^ and laborious to execute. Conjugation requires the creation and maintenance of a donor strain of *Escherichia coli*, which must contain a suitable broad-host-range conjugative plasmid in addition to the episome of interest; complications can arise if these two constructs share similar replication machinery, or if the episome of interest contains elements that are toxic when expressed in *E. coli*. The delivery of episomes through PEG transformation has been previously explored, but remains unutilized due to its low efficiency (≤ 1 transformant per 10^-8^ cells) and high variability between experiments^6^. Electroporation has a higher transformation efficiency (2.6 to 4.5 transformants per 10^5^ cells^5,7^), but has only been used to integrate non-replicative DNA into the nuclear genome. We sought to optimize the PEG and electroporation methods for episome delivery to *P. tricornutum,* with the hopes of developing a more rapid alternative to bacterial conjugation.

In this paper we describe optimized electroporation and PEG transformation protocols for engineering *P. tricornutum*. These methods are both faster and simpler to execute than bacterial conjugation, as well as more efficient than previously described protocols. We also demonstrate that it is possible to assemble episomes directly in the algal cell through non-homologous end joining (NHEJ) and homologous recombination (HR) pathways, forgoing the need for traditional *E. coli* or *Saccharomyces cerevisiae* cloning steps. Both protocols are suitable for rapid high throughput testing of several DNA constructs simultaneously and, due to their efficiency with low DNA amounts, could be used for the delivery of commercially synthesized episomes directly into the algal cell. Electroporation can also be used for the efficient delivery of Cas9 ribonucleoprotein (RNP) complexes, enabling DNA-free genetic engineering. The protoplasting techniques we established in *P. tricornutum* were used to successfully engineer *Thalassiosira pseudonana* cells via PEG transformation, demonstrating that these methods can be applied to other diatom species. These tools will be immensely valuable as the Pt-syn 1.0 project begins to build and test different synthetic chromosome designs.

## RESULTS

### Electroporation efficiency increases with enzymatic treatment of cells

Our initial attempts at electroporation using a previously described protocol^16^ that was adapted from the methods of Zhang and Hu^5^ yielded no transformants. After several failed attempts, we discovered that adjusting the electroporator capacitance from 25 to 50 µF yielded transformants for both episomes and marker cassettes, though efficiency was highly variable between experiments (Supplementary Table S1, Exp. 1 to 81). We used this method to deliver a linear 11 kb PCR-amplified episome, pPtGE31_ΔPtR (Addgene ID: 236260), to see if the construct would integrate into the genome or be replicated linearly in *P. tricornutum*. To our surprise, all analysed transformants demonstrated that the linear episome had been circularized in the algal nucleus through NHEJ at its termini.

When this electroporation protocol was repeated using cells that had reached the late stationary phase of growth, we obtained up to 272 transformants per 10^8^ cells with the same 11 kb PCR-amplified episome (Supplementary Table S1, Exp. 82 to 89). This represented a nearly 20-fold increase in efficiency when compared to transformations with early-log phase cultures (Exp. 80 and 81). Stationary phase cultures demonstrated noticeable differences in cell phenotype when compared to early-log phase cultures, with there being a higher abundance of oval-morphotype cells and protoplasts (i.e., diatoms lacking a cell wall) in the older cultures. *P. tricornutum* cells can appear in three different morphotypes – fusiform, oval, and triradiate – and can switch morphotypes in response to environmental stimuli^17,18^. This led us to hypothesize that morphotype and cell wall integrity were integral to electroporation transformation efficiency.

To test this hypothesis, we spheroplasted early-log phase *P. tricornutum* cells harvested from agar plates. Our previous experiments had used early log phase liquid cultures, which predominantly consisted of the fusiform morphotype; in past experiments, we had noticed a higher abundance of the oval morphotype in solid cultures harvested at the same growth stage. Plate-derived cells were spheroplasted using the enzyme alcalase, a serine protease, which has been previously shown to degrade the proteinaceous components of the *P. tricornutum* cell wall^19^. We observed that 100 µl of alcalase at an activity of approximately 3 Anson units per ml (i.e., 300 mAnson units) was enough to fully protoplast 3 x 10^8^ plate-derived cells when resuspended in 375 mM D-sorbitol (Fig. 1A and B). The protoplasted cells were very fragile and prone to rupture from osmotic pressure or physical damage; treatment of 3 x 10^8^ cells with 10 µl of alcalase (30 mAnson units) was enough to protoplast some cells, whilst leaving others with partially degraded cell walls (i.e., spheroplasts).

**Figure 1.**
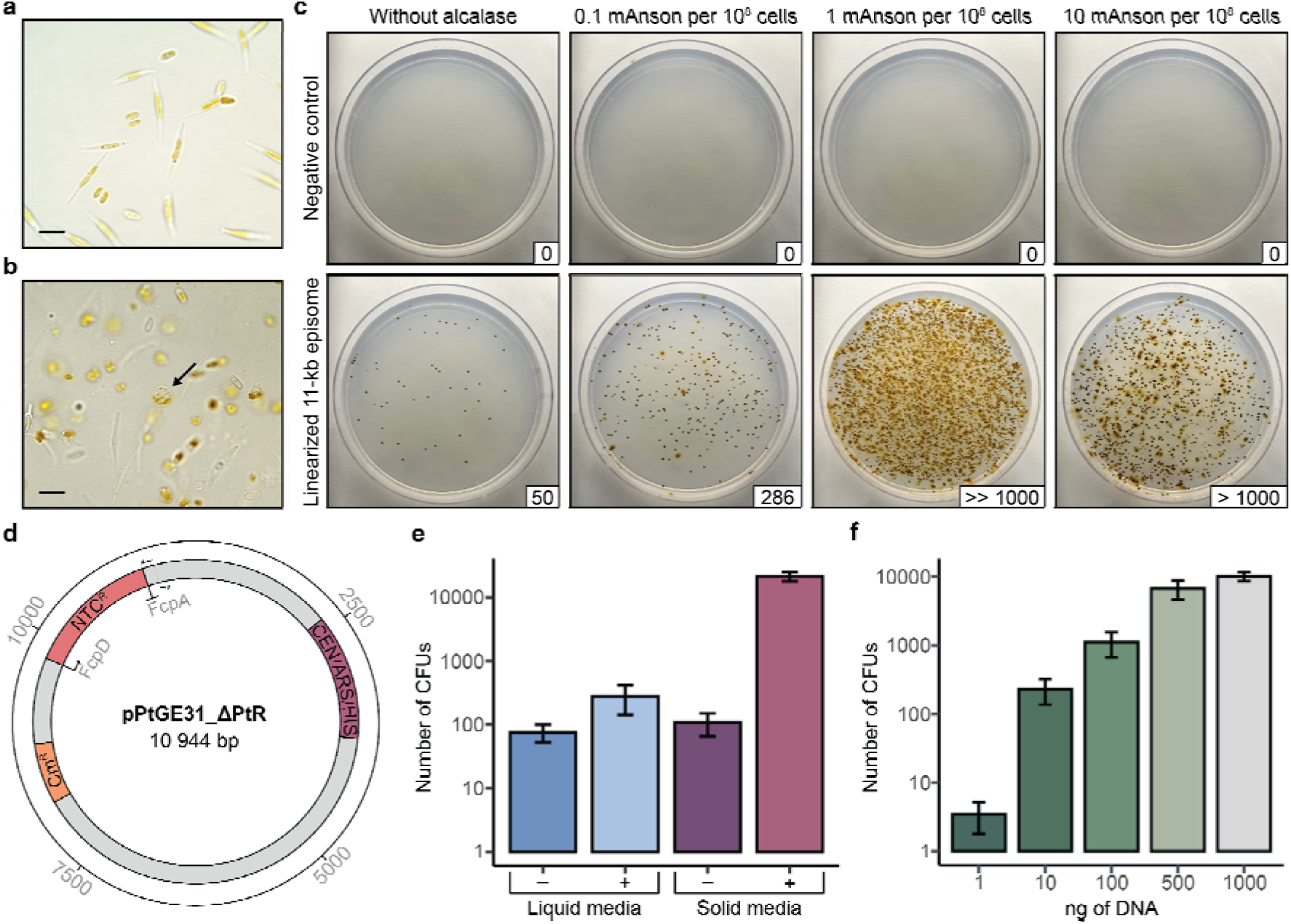
Spheroplasting *P. tricornutum* cells increases transformation efficiency. (**a**) Plate-derived *P. tricornutum* cells pre- and (**b**) post-treatment with 100 mAnson units of alcalase per 10^8^ cells. The arrow indicates a completely protoplasted cell, and the scale bars represent 10 µm. (**c**) Comparison of agar plate derived *P. tricornutum* cells treated with differing amounts of alcalase prior to electroporation. Cells were either electroporated with no DNA (negative control) or the 11 kb episome pPtGE31_ΔPtR, which was obtained through PCR amplification. The number of colony forming units (CFUs) is depicted in the bottom right corner for each transformation plate. Following electroporation, half of the total reaction was plated on ¼-salt L1 plates supplemented with 100 µg/ml nourseothricin. (**d**) A plasmid map of pPtGE31_ΔPtR, the episome used for transformations in panels c, e, and f. Primers depict where the plasmid was linearized during PCR-amplification. This plasmid contains a yeast centromere (CEN) and origin of replication (ARS) sequence, which enables episome replication and stability in *P. tricornutum*^6^, as well as a bacterial chloramphenicol resistance marker (Cm^R^) and an algal nourseothricin resistance marker (Ntc^R^) driven by the FcpD/FcpA promoter-terminator pair^9^. (**e**) A comparison of transformation efficiency for cells cultured in liquid or solid media with (+) or without (-) alcalase treatment prior to electroporation. (**f**) The number of *P. tricornutum* CFUs obtained with different nanograms of DNA during electroporation. Error bars denote the standard error of the mean from at least four biological replicates per experiment.

Plate-derived cells (3 x 10^8^) were treated with 0.3 mAnson units, 3 mAnson units, and 30 mAnson units of alcalase prior to the washing steps of the electroporation protocol. Transformation efficiency increased in all cases (Fig. 1C) when electroporating the same 11 kb PCR-amplified episome as before (Fig. 1D), with the 3 mAnson treatment demonstrating the highest increase in efficiency. Based on this result, we went forward with using 1 mAnson unit of alcalase per 10^8^ cells when spheroplasting *P. tricornutum* for electroporation.

### Electroporation efficiency of cells grown in liquid and solid media

We tested the electroporation efficiency of *P. tricornutum* cells grown in liquid media and on agar plates, with and without alcalase treatment, across five biological replicates (Fig. 1E) using the same PCR-amplified 11-kb episome as before. Though transformation efficiency varied between experiments, it was evident that the plate-derived alcalase-treated cells were drastically more efficient (∼1.2 transformants per 10^4^ cells) than untreated plate-derived cells (∼6.0 transformants per 10^7^ cells), untreated liquid-derived cells (∼4.2 transformants per 10^7^ cells), and treated liquid-derived cells (∼1.6 transformants per 10^6^ cells, Supplementary Table S2). Based on this result, all proceeding electroporations were conducted with cells derived from agar plates and spheroplasted with 1 mAnson unit of alcalase per 10^8^ cells. This optimized method was compared with the original and adjusted electroporation protocols to demonstrate the impact that changing the capacitance parameter and spheroplasting has on electroporation efficiency (Fig. S1).

### Transformation efficiency with varying amounts of DNA

Our prior transformations were all conducted with ∼500 ng of a PCR-amplified 11-kb plasmid, pPtGE31_ ΔPtR (Fig. 1D). We were interested to see how transformation efficiency varied when using differing amounts of this construct, and most importantly, what the minimum amount of DNA for transformation is. We electroporated as little as 1 ng and up to 1 µg of PCR-amplified pPtGE31_ ΔPtR across four biological replicates (Fig. 1F, data summarized in Supplementary Table S3). In all but one experiment, we were able to recover transformants when using just 1 ng of DNA during electroporation. Increasing the amount of DNA to 10 ng routinely gave rise to hundreds transformants, with there being an average of 372 CFUs per reaction. Interestingly, there was only a small change in transformation efficiency when the amount of DNA was doubled from 500 ng to 1 µg, suggesting that 500 ng of linear pPtGE31_ ΔPtR was enough to nearly saturate the electroporation reaction.

### Transforming linear and circular episomes to the algal nucleus

Our past explorations had relied upon the use of linear PCR-amplified DNA with blunt, non-phosphorylated termini. To test the efficiency of different structural forms of DNA, we prepared other variations of pPtGE31_ ΔPtR (Fig. 2A). The termini of the linear constructs varied in position along the episome and/or the properties of the termini (i.e., blunt or sticky ends, with or without 5′ phosphorylation). These constructs were adjusted to be equimolar prior to conducting electroporation to *P. tricornutum*. We found that there were nearly double the amount of CFUs when transforming any structural form of linear DNA compared to the circular episome (Fig. 2B).

**Figure 2.**
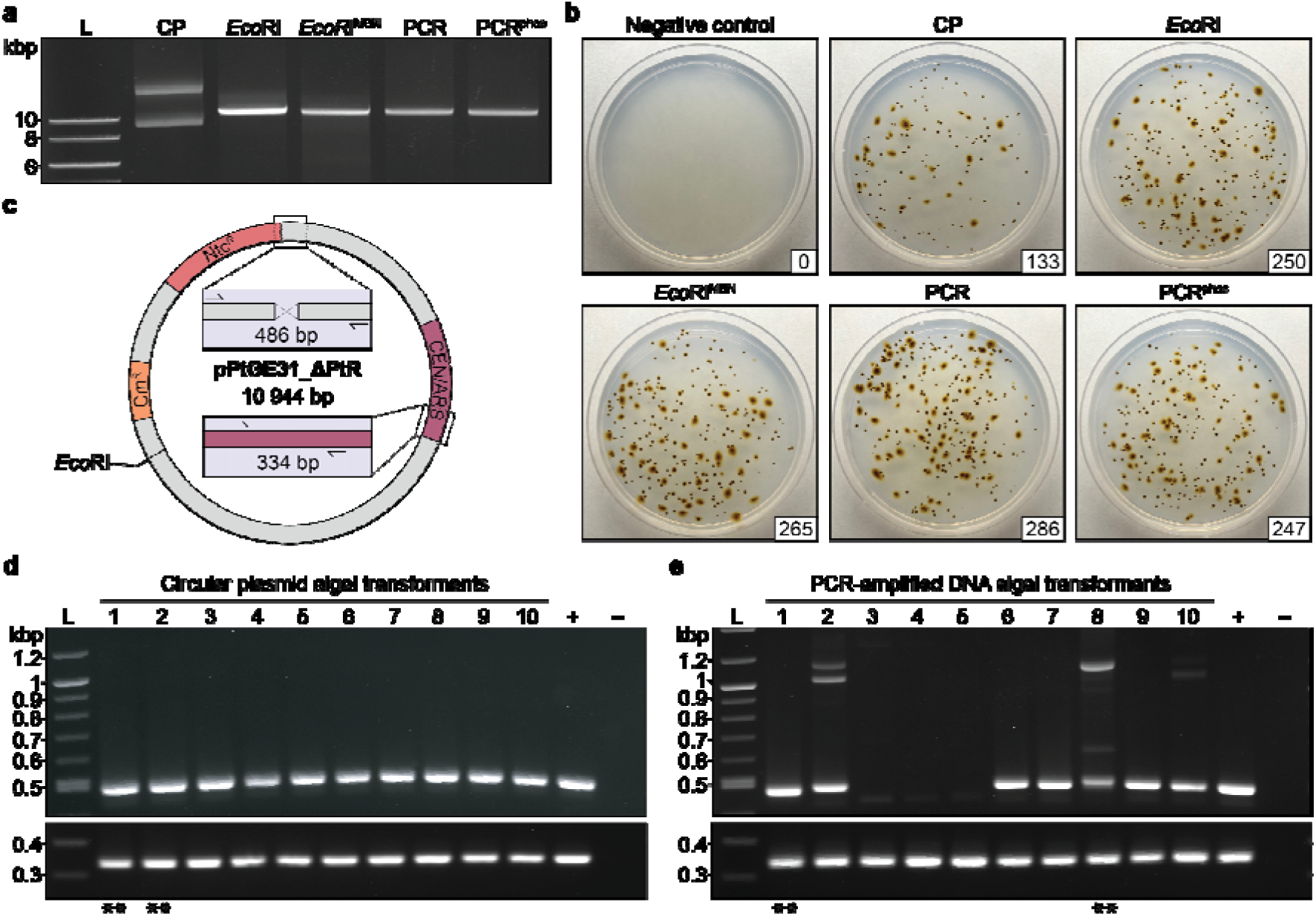
Comparing the transformation efficiency of linear, circularized, and phosphorylated DNA. (**a**) An agarose gel depicting different preparations of the episome pPtGE31_ ΔPtR. CP: circular plasmid, *Eco*RI: restriction enzyme digested plasmid, *Eco*RI^MBN^: restriction enzyme digested plasmid with additional mung bean nuclease treatment to remove sticky ends, PCR: PCR-amplified linear plasmid DNA, and PCR^phos^: PCR-amplified linear plasmid DNA generated with phosphorylated primers. (**b**) Comparison of *P. tricornutum* transformants electroporated with different preparations of pPtGE31_ ΔPtR. Following electroporation, one-tenth of the total reaction was plated on ¼-salt L1 plates supplemented with 100 µg/ml nourseothricin. Plates are labelled according to the type of DNA that was transformed, as described above. The number of CFUs is shown in the bottom right corner for each plate. (**c**) A plasmid map depicting the regions that were screened when assessing passaged algal transformants. The 486 bp junction spans the region where pPtGE31_ ΔPtR was split for PCR amplification, whereas the 334 bp junction spans a region of the plasmid backbone. (**d**) Screening 10 algal colonies that had been transformed with circular pPtGE31_ ΔPtR or (**e**) PCR-amplified pPtGE31_ ΔPtR. Dilute pPtGE31_ ΔPtR and genomic *P. tricornutum* DNA were used for the positive and negative controls, respectively. Double asterisks (**) indicate the algal episomes that would be recovered in *E. coli* and sent for whole plasmid sequencing.

We had also demonstrated that electroporated linear pPtGE31_ ΔPtR gets circularized in the *P. tricornutum* nucleus, but we had not yet performed comprehensive screening to determine if any indels were accruing during this repair process. To explore what occurs during termini fusion, ten colonies were repatched from the transformations with circular and PCR-amplified pPtGE31_ ΔPtR for additional screening. Colonies were screened using a set of primers that spans where the termini are expected to recombine for the PCR-amplified construct, as well as a region of the episome backbone (Fig. 2C). Colonies transformed with circular DNA did not demonstrate any unexpected insertions or deletions in the screened regions (Fig 2D), as expected. Whole plasmid sequencing of circular colonies 1 and 2 confirmed that the episomes had not accrued any mutations or indels during transformation.

Colonies transformed with PCR-amplified DNA appeared to have indels in the region where termini fusion occurs but not in the intact backbone region (Fig. 2E). We conducted Sanger sequencing of the termini region for colonies 1, 2, and 6 to 10, which confirmed that indels of varying sizes had accrued in this region for every colony screened (Supplementary Fig. S2). Whole plasmid sequencing of linear colonies 1 and 8 further confirmed the presence of indels in the termini region where the episome was being fused together, whereas all other regions of the episome had remained unchanged. Interestingly, the ∼500-bp insertions in colonies 2, 8, and 10 were not complimentary to any regions of the episome or algal genome. When these insertions were queried using BLASTn, they aligned to various regions of the salmon and/or trout genome accessions.

Single stranded salmon sperm (ssss) DNA is used as a carrier during electroporation, and to our surprise, we had discovered that it could be captured during NHEJ of the episome termini. We attempted electroporation without ssssDNA to see if this would impact transformation efficiency and it resulted in a 15-fold reduction in the number of CFUs (Supplementary Fig. 3), highlighting its importance as an additive for efficient transformation.

### Electroporation and recovery of a large plasmid

It has been demonstrated in other organisms that as episome size increases, electroporation efficiency decreases^20,21^. We sought to test the limits of electroporation in *P. tricornutum* by attempting transformation with pSC5, a 55.6 kb broad-host-range conjugative episome^22^ (Fig. 3C, Addgene ID: 188602).

**Figure 3.**
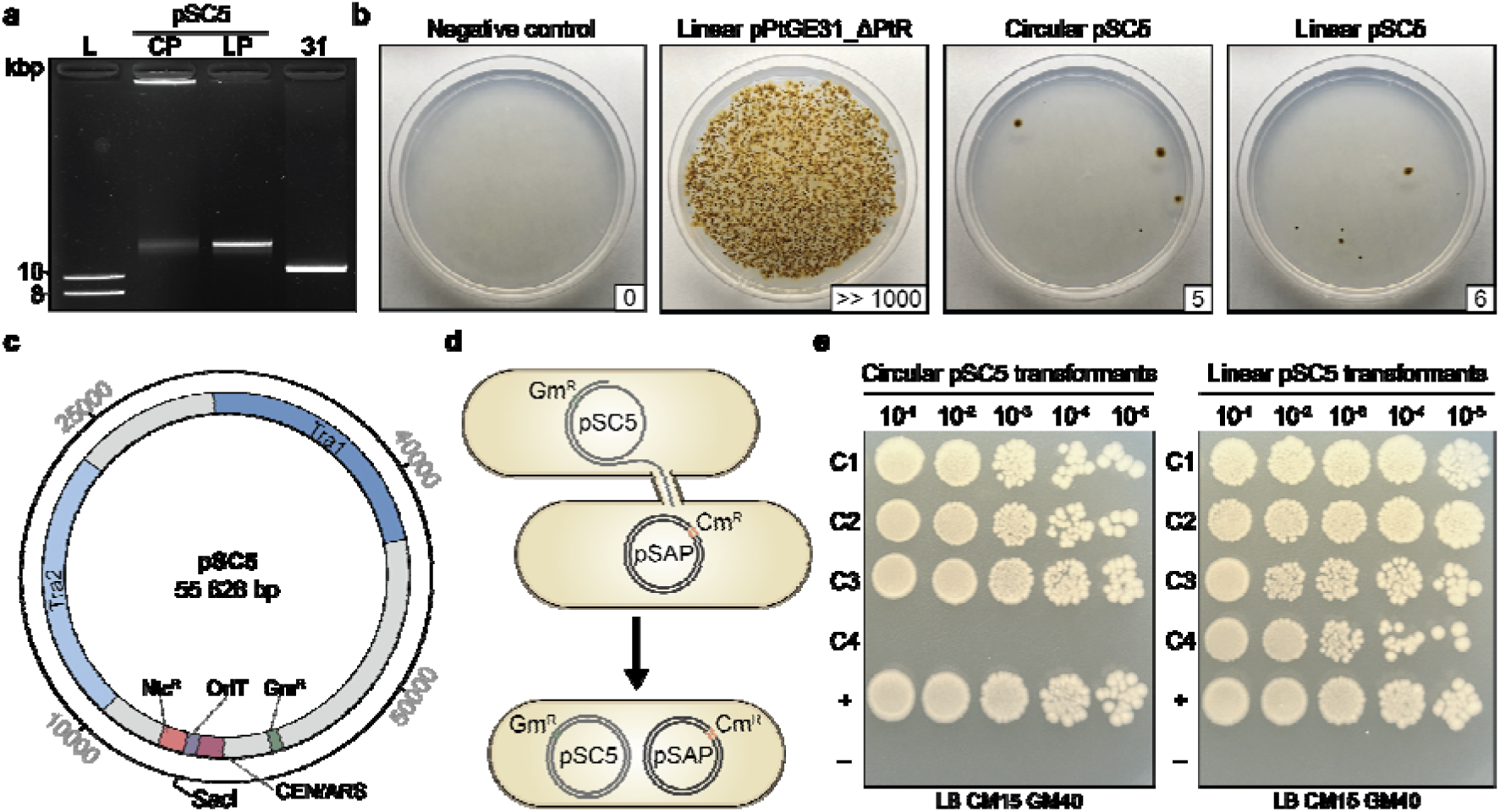
Electroporation and recovery of a 55.6 kb conjugative plasmid. (**a**) An agarose gel depicting the circular pSC5 (CP), linearized pSC5 (LP), and PCR-amplified pPtGE31_ ΔPtR (31). (**b**) Electroporation of circular or linear variants of pSC5 and PCR-amplified pPtGE31_ ΔPtR into *P. tricornutum* cells. Following electroporation, half of the total reaction was plated on ¼-salt L1 plates supplemented with 100 µg/ml nourseothricin. Plates are labelled according to the type of DNA that was transformed, as described above. The number of CFUs is shown in the bottom right corner for each plate. (**c**) A plasmid map of pSC5. The transfer regions (Tra1 and Tra2) encode for essential conjugative proteins. The origin of transfer (OriT) serves as a recognition sequence for the conjugative protein apparatus, facilitating the transfer of pSC5 during conjugation. The plasmid contains selection markers for bacteria (Gm^R^, gentamycin resistance) and *P. tricornutum* (Ntc^R^, nourseothricin resistance) as well as a yeast centromere (CEN) and replication origin (ARS), which enable episome stability and replication in *P. tricornutum*. (**d**) A donor strain of *E. coli* harbouring pSC5, which confers gentamycin resistance, conjugating to a recipient strain of *E. coli* harbouring pSAP, which confers chloramphenicol resistance. Successful conjugation leads to the creation of a transconjugant strain that is gentamycin and chloramphenicol resistant. (**e**) Testing the conjugation efficiency of *E. coli* strains harbouring episomes that had been recovered from *P. tricornutum* transformants electroporated with circular or linear pSC5. Strains harbouring pSC5 or pSAP were used as the positive and negative controls, respectively. Dilutions of 10^-1^ to 10^-5^ were spot plated on LB plates supplemented with 15 µg/ml chloramphenicol and 40 µg/ml gentamicin.

Equimolar amounts of circular and linear pSC5 were prepared along with PCR-amplified pPtGE31_ ΔPtR (Fig. 3A). Electroporation with either form of pSC5 gave rise to transformants, though the number of CFUs was orders of magnitude lower than electroporation with a linear 11 kb episome (Fig. 3B), as anticipated. Transformation efficiency with linear pSC5 was three times greater than with the circular form when averaged across three biological replicates (Supplementary Table S4), with there being an average of 12 CFUs per reaction.

To determine if the complete 55.6 kb episome had been successfully electroporated into *P. tricornutum*, we passaged four colonies from each of the transformations with circular and linear pSC5. DNA was isolated from the algal colonies and electroporated into *E. coli*. Then, we performed conjugation using a pool of *E. coli* colonies for each transformation (Fig. 3D). Here, if the transformants contain an intact version of pSC5, the episome will facilitate conjugation and self-mobilization into a recipient strain harbouring pSAP^12^ (Addgene ID: 206429), a 10 kb plasmid that contains a chloramphenicol resistance marker. Successful transconjugants will harbour pSC5 and pSAP, therefore carrying resistance to both gentamycin and chloramphenicol. Conjugation was successful for seven of the eight *E. coli* pools, demonstrating that intact pSC5 had been successfully electroporated and maintained in the majority of screened *P. tricornutum* transformants (Fig. 3E).

### Electroporation and recombination of multiple DNA fragments

#### i. Assembly via the NHEJ pathway

The surprising discovery of salmon DNA inserted into the termini region of PCR-amplified pPtGE31_ΔPtR led us to hypothesize that *P. tricornutum* could assemble multiple DNA fragments into a single episome through NHEJ. To test this, we created an episome, pPtGE31_ShBle (Addgene ID: 236261), that contains the NAT and ShBle markers conferring resistance to nourseothricin and zeocin, respectively. The episome pPtGE31_ShBle was then amplified as two non-overlapping fragments, each containing one of the algal selective markers. The fragments were adjusted to be equimolar prior to electroporation into *P. tricornutum* (Fig. 4A).

**Figure 4.**
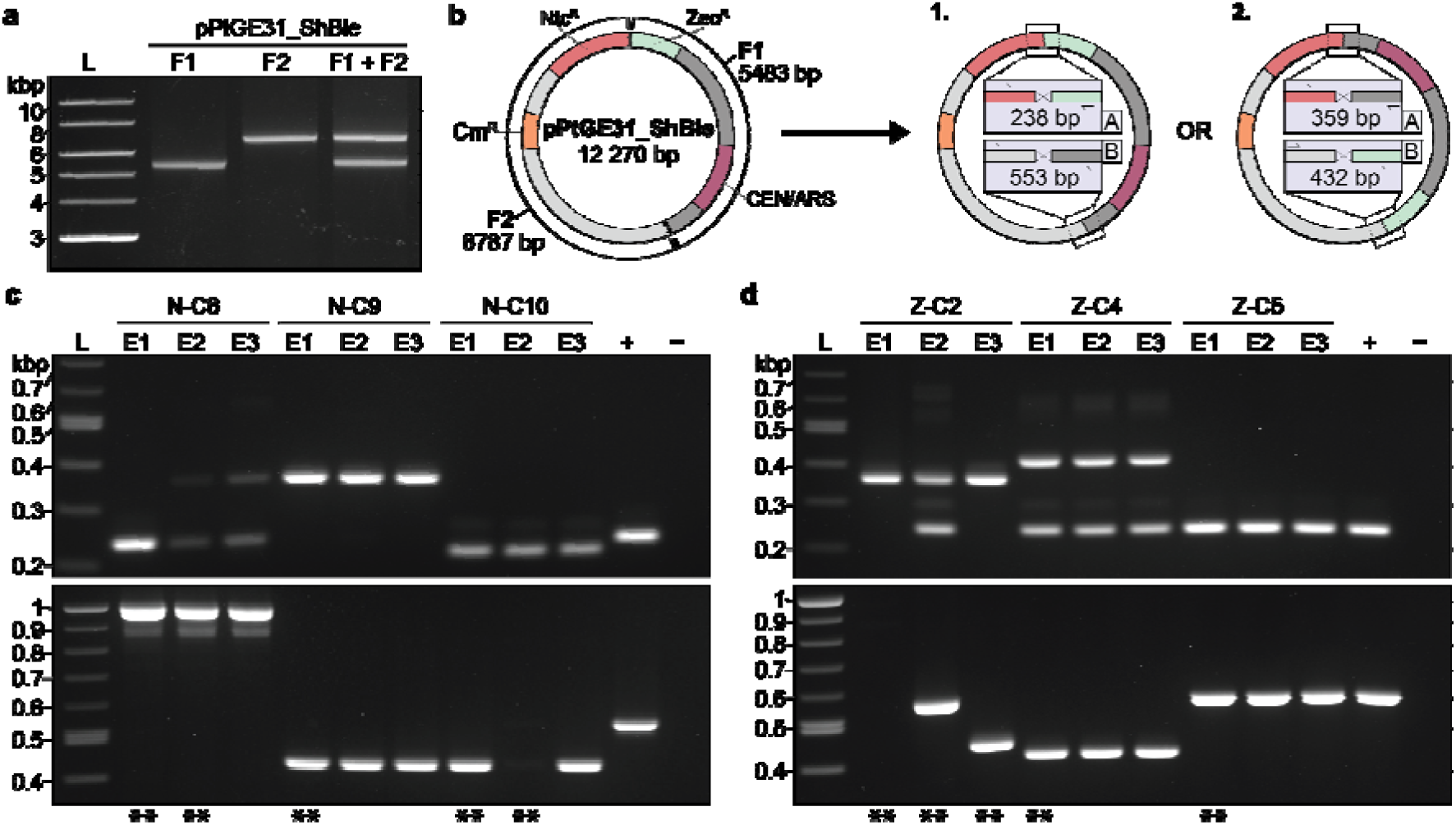
Electroporation and assembly of two non-overlapping fragments in *P. tricornutum*. (**a**) The episome pPtGE31_ShBle was amplified as two fragments (F1: fragment 1, F2: fragment 2). The fragments were adjusted to be equimolar before combining them into an assembly mixture (F1 + F2). (**b**) A plasmid map of pPtGE31_ShBle and a depiction of the different possible orientations post-assembly. The episome contains algal selection markers for Ntc^R^ and Zeo^R^, a bacterial Cm^R^ selection marker, as well as a yeast CEN and ARS sequences. F1 and F2 do not share any overlapping sequences and thus can assemble in two different orientations. Primer sets were designed to span the two junctions, A and B, where recombination is expected to occur. The amplicons will vary in size depending on the orientation of the fragments. (**c**) Screening three *E. coli* transformants per algal colony using primers spanning junctions A and B. DNA were isolated from algal transformants that were electroporated with F1 + F2 and selected initially with nourseothricin or (**d**) zeocin. Dilute pPtGE31_ ShBle and water were used for the positive and negative controls, respectively. Double asterisks (**) indicate the *E. coli* colonies that were sent for whole plasmid sequencing.

Cells transformed with both fragments simultaneously were plated on nourseothricin- or zeocin-containing plates initially (Supplementary Fig. S4). Out of the 116 colonies passaged post-electroporation, 94 demonstrated double resistance to both nourseothricin and zeocin (Supplementary Table S5). Twenty transformants, ten derived from each initial transformation plate, were passaged onto double-selection media, then DNA isolation and PCR screens were performed. These colonies are denoted with an N- or Z- to depict which initial selection plate they were derived from (N: nourseothricin, Z: zeocin).

Two orientations are expected when the fragments amplified from pPtGE31_ShBle are re-assembled (Fig. 4B). We designed a PCR screen that would not only assess if both fragments had been recombined, but could also determine the orientation of the fragments. When screening algal transformants, we only used the primer set spanning junction B as junction A contains primers that bind to the FcpD promoter and FcpA terminator, which are present in the episome and algal genome. Of the twenty colonies screened, nine had amplification across junction B, with only five colonies demonstrating the expected amplicon sizes (Supplementary Fig. S5). The other four colonies either demonstrated multiple amplicons, some of which were the expected sizes, and/or amplicons that were larger than expected. Based on this initial screen, DNA from colonies N-C6, N-C9, N-C10, Z-C2, Z-C4, and Z-C5 were transformed into *E. coli* for further screening.

Three *E. coli* transformants were assessed per algal colony using the primer sets spanning junctions A and B (Fig. 4C and D). Interestingly, the *E. coli* transformants derived from algal colonies N-C6, N-C10, and Z-C2 demonstrated different banding patterns, suggesting that these algal colonies likely had a pool of assembled episomes. All *E. coli* transformants from algal colony N-C9 depicted the expected banding pattern for orientation 2, whereas all *E. coli* transformants from algal colony Z-C5 depicted the expected banding pattern for orientation 1. DNA from *E. coli* colonies N-C6-E1/E2, N-C9-E1, N-C10-E1/E2, Z-C2-E1/E2/E2, Z-C4-E1, and Z-C5-E1 were sent for whole-plasmid sequencing to validate if assembly had occurred.

The sequencing results demonstrated that recombination of fragments 1 and 2 had occurred in all assessed *P. tricornutum* transformants (Supplementary Fig. S6). In the majority of assembled episomes, multiple partial copies of fragment 1 and 2 had recombined. One extreme case was *E. coli* colony Z-C2-E2, which harboured a 41.5 kb episome containing eight partial or whole copies of both fragments 1 and 2. Only *E. coli* colonies N-C9-E1, N-C10-E1, Z-C2-E3, and Z-C5-E1 contained the expected assembled episomes. Sequencing of *E. coli* transformants derived from the same algal colony revealed that algal colonies N-C6, N-C10, and Z-C2 possessed a pool of assembled episomes, as we had hypothesized.

### ii. Assembly via the HR pathway

Though it is possible to assemble episomes via the NHEJ pathway, screening and sequencing demonstrated that the majority of *P. tricornutum* transformants contained undesirable assemblies of the non-overlapping fragments. To be able to reliably assemble episomes in the algal nucleus, an alternative strategy is needed. We sought to test whether overlapping fragments could be simultaneously transformed and assembled through the HR repair pathway. It was hypothesized that the overlapping sequences between fragments would enable site-specific recombination at the termini, thereby creating episomes of an expected size and directionality.

To test this hypothesis, we split the episome pPtGE31_ShBle into two, three, or four fragments that overlapped with each other by ∼160 bp (Fig. 5A). Fragments were generated by PCR amplification using PtGE31_ΔPtR (Addgene ID: 236260) or pPtGE31_ShBle as template DNA (Table S7). In all assembly designs, one of the overlaps was positioned within the open reading frame (ORF) for nourseothricin N-acetyl transferase (NAT), which confers nourseothricin resistance. Thus, when the assembly fragments are electroporated, nourseothricin-resistant transformants should only appear if homologous recombination has occurred within the NAT ORF or if there is carry-over of the PCR template DNA. Primers were designed to screen the other overlapping termini for successful recombination (Fig. 5A, bottom boxes).

**Figure 5.**
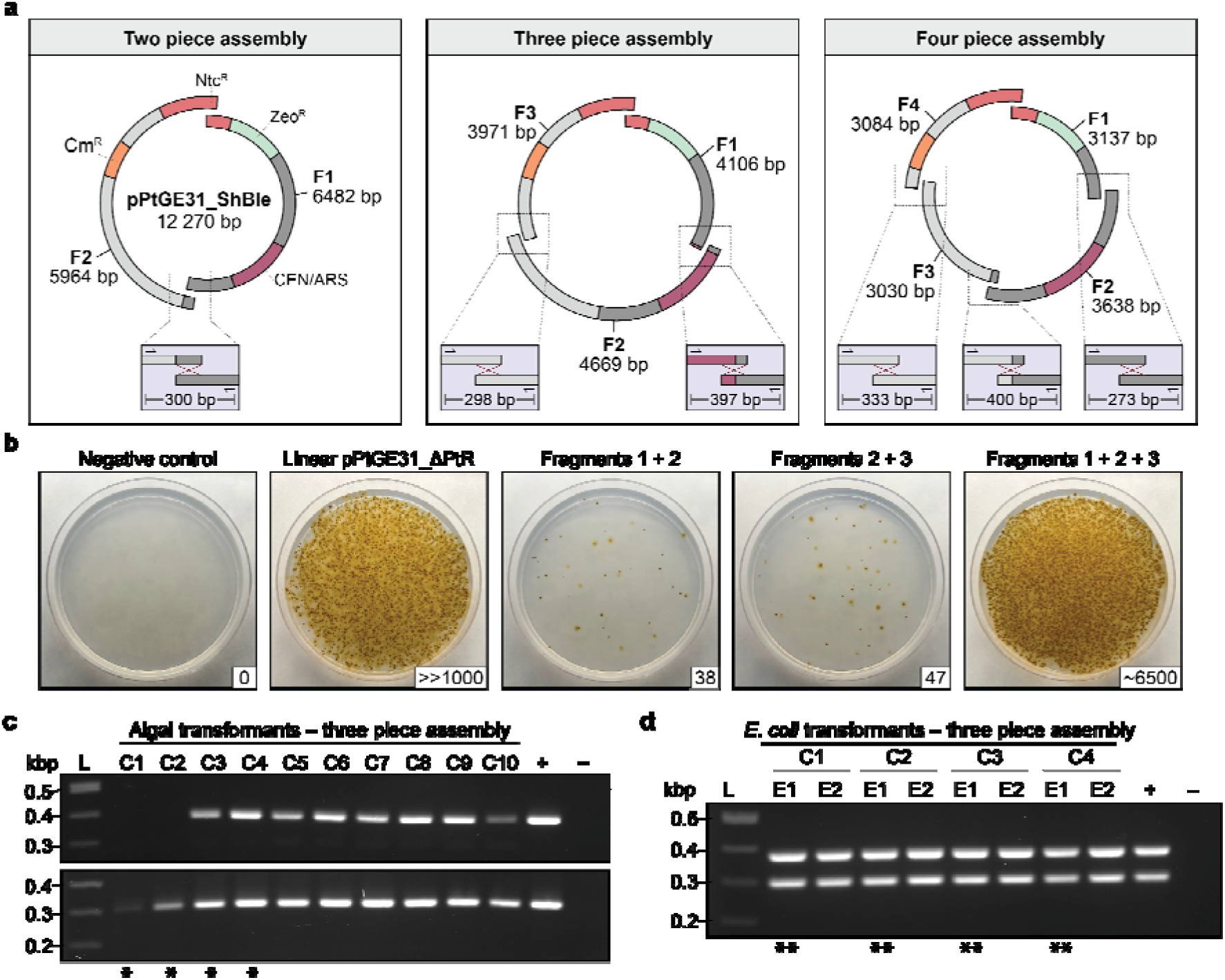
Electroporation and assembly of overlapping fragments in *P. tricornutum*. (**a**) The episome pPtGE31_ShBle was amplified as two, three, or four overlapping fragments (F1: fragment 1, F2: fragment 2, F3: fragment 3, F4: fragment 4). The fragments were adjusted to be equimolar before combining them into their respective assembly mixtures. Primers were designed to screen the other overlapping junction(s), demonstrated in the bottom boxes. The episome contains a bacterial Cm^R^ marker, the algal Ntc^R^/Zeo^R^ markers, and a yeast CEN/ARS. (**b**) Transformation of three overlapping fragments via electroporation. Linear pPtGE31_ ΔPtR (500 ng) was used as a positive control. The fragments were also electroporated in combinations (i.e., <u>F1 + F2</u> or <u>F2 + F3</u>) to determine the rate of carry-over of the template episome. For all transformations, half of the total reaction was plated on ¼-salt L1 plates supplemented with 100 µg/ml nourseothricin. The number of CFUs is shown in the bottom right corner for each plate. (**c**) Screening 10 algal transformants post electroporation with three overlapping fragments and (**d**) eight *E. coli* transformants with episomes recovered from C1, C2, C3, and C4. The junctions screened span F1/F2 and F2/F3. For the algal screen, the positive control consists of dilute pPtGE31_ShBle and the negative control consists of wild-type *P. tricornutum* DNA. The same positive control was used for E. coli screens, however, the negative control consisted of water. Single asterisks (*) indicate the algal colonies that had their DNA isolated and transformed into *E. coli*. Double asterisks (**) indicate the *E. coli* colonies that were sent for whole plasmid sequencing.

We first tested the two-piece HR assembly by electroporating fragments 1 and 2 individually or simultaneously (Fig. S7A). When electroporated individually, fragments 1 and 2 gave rise to 8 and 2 CFUs, respectively, indicating that carry-over of the PCR template DNA had occurred. However, when the fragments were electroporated simultaneously, there were more than 1800 CFUs – far more than what would be anticipated by carry-over alone. Ten algal colonies were selected for screening of the other fragment junction (Fig. S7B), all of which demonstrated the expected 300 bp band, suggesting that successful homologous recombination had occurred. Episomes from algal colonies C1, C2, and C3 were recovered in *E. coli*. Three *E. coli* colonies were screened per initial algal colony, all of which demonstrated the correctly sized band (Fig. S6C). To confirm if the fragments had been correctly recombined, episomes from *E. coli* colonies C1-E1, C2-E1, and C3-E1 were sent for whole plasmid sequencing. This demonstrated that colonies C1-E1 and C3-E1 contained the correctly assembled episome (Fig. S8), but the episome recovered from C2-E1 had two partial copies of fragment 1.

The same methodological approach was used to electroporate three overlapping fragments simultaneously (Fig. 5B). Electroporation of fragments 1 and 2 or 2 and 3 gave rise to 38 and 47 CFUs respectively, whereas electroporation of all three fragments gave rise to ∼6500 CFUs. Ten algal colonies were screened for the junctions between F1/F2 or F2/F3. All colonies contained the expected bands except for C1 and C2, which lacked an amplicon for the F1/F2 junction and demonstrated a weak amplicon for the F2/F3 junction (Fig. 5C). We anticipated that assembly may have failed in these colonies, but to further investigate this, we attempted to recover the algal episomes from C1 and C2 in *E. coli*, along with the positively screened episomes from C3 and C4. We were able to recover *E. coli* transformants for all the tested episomes and decided to screen two CFUs per initial algal colony, all of which demonstrated correctly sized amplicons for both junctions (Fig. 5D). Episomes isolated from *E. coli* colonies C1-E1, C2-E1, C3-E1, and C4-E1 were sent for whole plasmid sequencing, with only C1-E1 deviating from the expected assembly (Fig. S8).

To further test the limits of the HR assembly pathway, we attempted electroporation of four overlapping fragments simultaneously. Electroporation of only three of the four fragments (i.e., <u>F1 + F2 + F3</u> or <u>F2 + F3 + F4</u>) gave rise to 85 and 56 transformants, demonstrating the carry-over of template DNA once again (Fig. S7D). However, when all four fragments were combined and transformed simultaneously, there was a >20-fold increase in the number of CFUs. Ten algal colonies were screened across the remaining three junctions, and all but C1 and C7 demonstrated the expected amplicons (Fig. S7E). Colonies C1 and C7 specifically lacked the 273 bp amplicon for the junction between F1/F2. Algal episomes from C1, C2, C3, and C7 were transformed into *E. coli*; however, C7 did not give rise to any bacterial transformants, suggesting that incomplete assembly had occurred. Two *E. coli* colonies were screened for the episomes derived from C1, C2, and C3, all of which demonstrated the expected amplicons for the three junctions. Whole plasmid sequencing of C1-E1, C2-E1, and C3-E1 revealed that correct assembly had occurred in algal colonies C1 and C3 (Fig. S7), but C2 contained mutliple partial copies of fragments 1 and 4.

In total, 26 of the 30 algal colonies screened across the various assemblies demonstrated that recombination had successfully occurred across the tested junctions. From the episomes recovered in *E. coli*, 7 out of 10 had the correctly assembled episomes when fully sequenced. This demonstrates that it is both feasible and reliable to assemble episomes through the HR pathway in *P. tricornutum*.

### Delivery of RNPs via the optimized electroporation protocol

RNP complexes can be delivered to *P. tricornutum* through biolistic bombardment^23^, providing a means for DNA-free genome engineering. We anticipated that it would also be possible to electroporate RNPs into this species, but this had yet to be reported in any diatoms. As a proof-of-concept, we replicated the methodology Serif et al. had used to deliver RNPs^23^ using our optimized electroporation protocol in lieu of biolistics.

Serif et al. had previously shown that knock-out of the *P. tricornutum Adenine Phosphoribosyl Transferase* (*PtAPT*) gene confers resistance to the cytotoxin 2-fluoroadenine (2-FA) at a dose of 10 µM^23^. The locus was targeted using two guide RNAs (gRNAs), each of which allowed for biallelic knock-out^23^; we opted to use the first gRNA (gAPT1) for our proof-of-concept experiments as it had generated 11 of the 16 2-FA resistant transformants in their singleplex experiments. HiFi Cas9 and a single guide RNA for APT1 (sgAPT1) were complexed according to the manufacturer’s guidelines (Fig. 6A). Then, the RNP complex was combined with ssssDNA and electroporated into spheroplasted *P. tricornutum* cells. We opted to include carrier DNA when performing the RNP experiments as ssssDNA is thought to bind to and neutralize the positive charge of the cell membrane during electroporation, making it easier for other charged biomolecules to pass through^24^.

**Figure 6.**
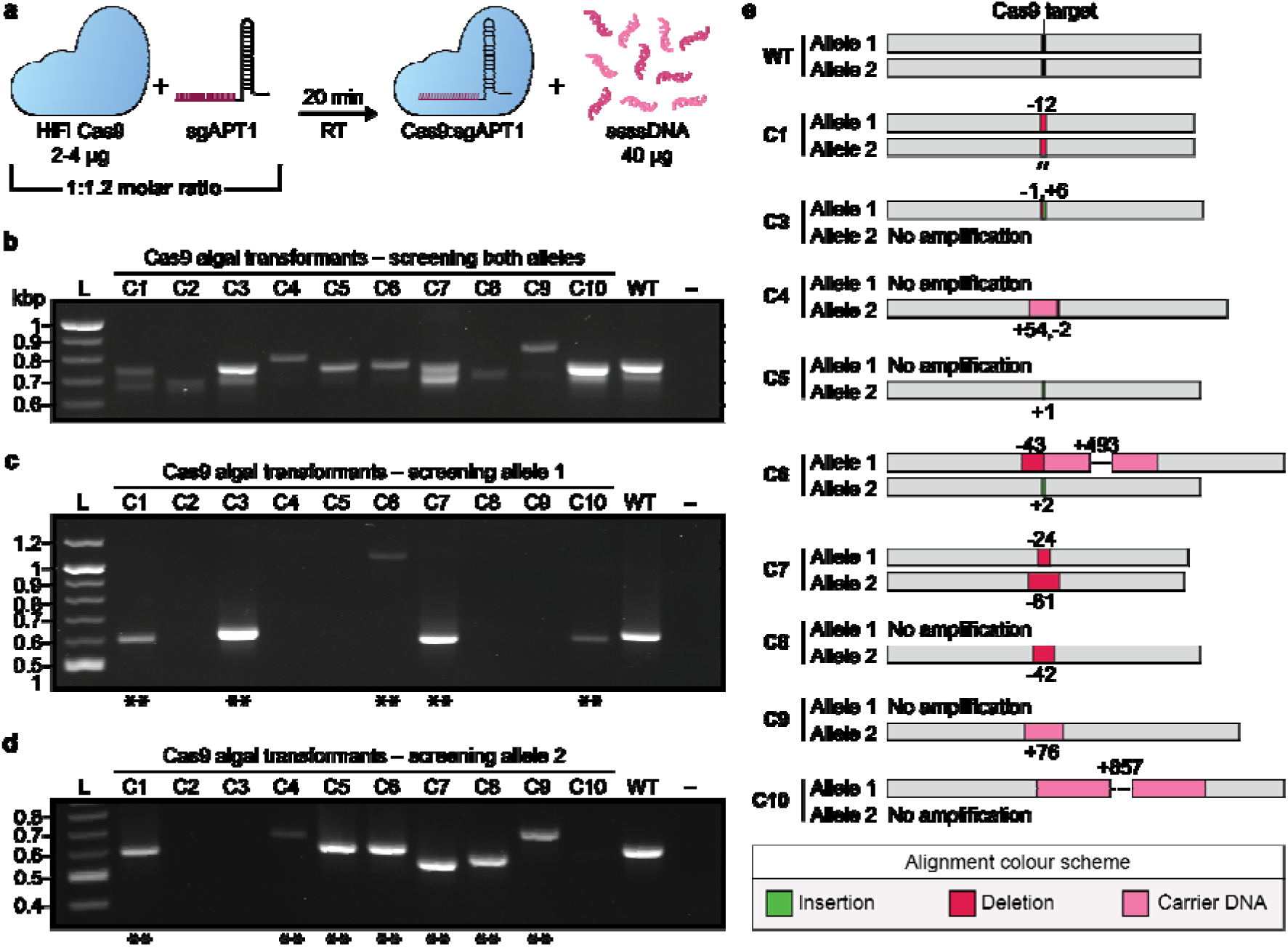
Electroporation of a Cas9:sgAPT1 complex into *P. tricornutum* for DNA free engineering. (**a**) The HiFi Cas9 complex is combined with a sgRNA that targets the *PtAPT* locus (sgAPT1). After twenty minutes at room temperature, the complex is combined with 40 µg of carrier DNA (ssssDNA) prior to electroporation. (**b**) Screening ten 2-FA resistant algal transformants with primers that bind to both *PtAPT* alleles, (**c**) allele 1, and (**d**) allele 2. The positive and negative controls consist of wild-type *P. tricornutum* DNA and water, respectively, for all PCR screens. Double asterisks (**) indicate the amplicons that were sent for Sanger sequencing. (**e**) Alignment of the Sanger sequences captured for algal colonies C1 and C3-C10.

Transformants were recovered for one day before plating one-fifth of the reaction across ½ salt L1 plates supplemented with 10 µM or 5 µM 2-FA, with or without 5 µg/ml adenine (Fig. S9). This pilot experiment generated thousands of 2-FA resistant transformants across the plates that had been supplemented with adenine. Ten 2-FA resistant colonies were passaged for further screening at the *PtAPT* loci to determine if Cas9-mediated engineering had occurred.

We used the primer sequences reported by Serif et al.^23^ (Table S7) to screen the *PtAPT* alleles using PCR. When using primers that bind to both alleles, amplicons of varying sizes were visible across the 2-FA resistant transformants, most of which differed from that of the wild-type strain (Fig. 6B). Likewise, amplicons of varying sizes were visible when using primers specific to allele 1 (Fig. 6C) and 2 (Fig. 6D). This suggested that Cas9-mediated engineering had occurred, but to corroborate our findings, amplicons generated with allele-specific primers were sent for Sanger sequencing.

Alignment of the sequenced amplicons to the wild-type sequence for alleles 1 and 2 confirmed the presence of indels in all 12 alleles that could be analysed (Fig. 6E). We were able to generate sequencing data of both alleles for three of the transformants (C1, C6, and C7), with one clone (C1) demonstrating a homozygous 12 bp deletion across both alleles. Interestingly, there were large (> 50 bp) insertions in four of the sequenced alleles. Querying the insertion sequences in NCBI blast revealed that carrier DNA had integrated into these loci during the double-strand break repair process. The alignments for C6 and C10 contained two and four non-contiguous regions from the salmon and/or trout genomes, demonstrating that multiple shorn pieces of carrier DNA had been integrated as opposed to a single large fragment.

### Cotransformation of an RNP complex and episome

With the discovery that RNP complexes can be electroporated into *P. tricornutum*, we next sought to see if both an RNP and episome could be co-transformed simultaneously, and if so, what the rate of co-transformation is. To test this, the PCR-amplified episome pPtGE31_ΔPtR was combined with the Cas9:sgAPT1 complex and electroporated into spheroplasted *P. tricornutum* cells. Transformants were first plated on nourseothricin or 2-FA selective plates, and then later repatched onto the alternative selection type to determine if co-transformation had occurred.

When testing the sensitivity of wild-type *P. tricornutum* to varying 2-FA concentrations (0, 2.5, 5, 10, 15, 20 µM 2-FA) on ½-salt L1 plates supplemented with adenine (5 µg/ml), we noticed an absence of growth on the 0 µM 2-FA plate. This suggested that the presence of adenine alone could impact wild-type *P. tricornutum* growth. We tested this theory by plating cells onto ½-salt L1 plates supplemented with varying levels of adenine (0, 1, 2.5, 5 µg/ml; Figure S10). The growth of wild-type algae was negligible after one-week on the 5 µg/ml adenine plate, but relatively unimpacted on the 0.5 and 1 µg/ml plates. Relatedly, our earlier RNP transformation experiment (Fig. S9) had demonstrated that adenine supplementation is important when selecting for *PtAPT* knock-out colonies post electroporation. Thus, when co-transforming pPtGE31_ΔPtR and Cas9:sgAPT1, the electroporated cells were plated on two different selection plates: (1) ½-salt L1 plates supplemented with 5 µg/ml adenine and 10 µM 2-FA, and (2) ½-salt L1 plates supplemented with 1 µg/ml adenine and 100 µg/ml nourseothricin. The lower adenine concentration was chosen to allow for the growth of co-transformed colonies whilst not interfering with the growth of colonies that had only uptaken the episome.

After two weeks of growth on nourseothricin or 2-FA selection, 50 co-transformed colonies were repatched onto plates with the same selection type and grown for 1 week. Then, the colonies were repatched onto the alternative selection type to determine if any had double-resistance. Of the 100 screened colonies, 13 demonstrated double-resistance to nourseothricin and 2-FA (Table S6), indicating co-transformation had successfully occurred at a rate of ∼10%.

### Revisiting the PEG transformation method

A polyethylene glycol (PEG) transformation method for *P. tricornutum* with an efficiency of ≤ 1 transformant per 10^8^ cells was previously described by Karas et al^6^. For this method, algal cells were spheroplasted using a combination of zymolyase, lysozyme, and hemicellulase ahead of PEG treatment. We sought to revisit this method using alcalase as the sole enzyme for cell wall removal.

We followed the method described by Karas et al.^6^, substituting the aforementioned enzyme cocktail with 10, 100, or 1000 µl of alcalase for protoplasting ∼3 x 10^8^ cells. As well, the protoplasts were transformed with 500 ng of linear pPtGE31_ΔPtR as opposed to 25 µg of a circular episome. Treatment with 1000 µl of alcalase generated approximately 160 transformants per 3 x 10^8^ cells, an efficiency of ∼4 transformants per 10^7^ cells. The other two treatments yielded 0 transformants (10 µl of alcalase) and approximately 6 transformants (100 µl alcalase), so all future PEG transformations were conducted with 1000 µl of alcalase per 3-5 x 10^8^ cells.

We then sought to test other parameters of the PEG method to further optimize the transformation efficiency. First, we began to include 40 µg of carrier DNA per PEG reaction as this had led to a ∼15-fold increase in transformation efficiency during electroporation. Then, we tested various concentrations of PEG-8000 and found that a range from 20 to 30% (w/v) can be used (Figure S11A). Going forward with 20% PEG, we then tested if the duration of protoplasting, inclusion of carrier DNA, and length and/or temperature of incubation would alter transformation efficiency (Fig S11B). As anticipated, the inclusion of carrier DNA increased transformation efficiency >10-fold (Fig. S11C). Longer protoplasting and incubation times also appeared to increase transformation efficiency to ∼6 transformants per 10^6^ cells (Fig. S11C).

The optimized PEG transformation protocol was then used to transform 1 µg of pSC5 into protoplasted *P. tricornutum* cells, generating ∼60 transformants (Fig. S2D). DNA were isolated from four algal transformants and recovered in EPI300 *E. coli*. Bacterial transformants were then assessed for conjugation in the same manner as before. All recovered episomes demonstrated the ability to conjugate (Fig. S11E), verifying that pSC5 had been intactly transformed into *P. tricornutum* through the PEG method.

### PEG transformation to T. pseudonana

We performed PEG transformation to *T. pseudonana* using the optimized conditions established with *P. tricornutum*. Given the differences in cell wall composition between these species, *T. pseudonana* cells were subjected to varying protoplasting conditions (Fig. S12A) before PEG treatment. The circular episome pBIG1 conferring nourseothricin resistance and carrying eGFP was used in these experiments. All protoplasting conditions led to some degree of growth on the nourseothricin selection plates, with harsher protoplasting conditions (i.e., ≥ 500 µl alcalase) yielding more single colony growth.

Transformants from the 500 µl alcalase treatment plate were pooled and subjected to fluorescence microscopy to assess if eGFP was being expressed, as this would indicate successful episome transfer. Both autofluorescence and eGFP signals were visible in the pooled cells, demonstrating that episome transfer had occurred.

To determine if protoplasting was truly occurring, *T. pseudonana* cells were incubated with 10 or 1000 µl of alcalase for 40 minutes (Fig. S12B and C, respectively) and then imaged via microscopy. A higher proportion of protoplasted cells are visible when using a greater amount of the enzyme, as anticipated if protoplasting were occurring.

## DISCUSSION

In this paper, we demonstrate that partial or complete removal of the cell wall greatly increases the efficiency of electroporation and PEG transformation in *P. tricornutum*. These optimized protocols enabled us to push the limits and possibilities of transformation in diatoms. Episomes as large as 55.6 kb could be intactly delivered through either approach, and it was possible to electroporate RNP complexes directly to the algal nucleus for DNA-free genome engineering. Perhaps most interestingly, we also discovered that episomes can be directly assembled in the diatom cell with high efficiency – a feat that has been predominantly limited to *E. coli* and *Saccharomyces cerevisiae*, the workhorses of synthetic biology. These methods are not only more efficient than what has been reported in the past^5–7^, but present as faster and simpler alternatives to bacterial conjugation for the reliable transformation of episomes.

The cell wall is the first physical barrier to entry when attempting to transform diatoms. It follows that permeabilizing this wall would increase the ability for biomolecules, like DNA and RNPs, to successfully enter the cell. Spheroplasting and protoplasting techniques are commonly used for transforming other algal and plant species^25,26^, but had yet to be fully investigated in diatoms until this study. We found that spheroplasting *P. tricornutum* cells with alcalase prior to conducting electroporation increased the efficiency by ∼76-to 280-fold (Supplementary Table S2), with cells derived from plated cultures demonstrating the highest transformation rates. Protoplasting the cells also enabled efficient transformation through the PEG method. Though *P. tricornutum* is unique in both its poorly silicified cell wall^27^ and willingness to grow on solid media, we were able to adapt the alcalase-based protoplasting technique for *T. pseudonana*, a centric diatom species with a silicified cell wall. We believe that the spheroplasting and protoplasting techniques can be applied to other diatom species, though it will likely be necessary to optimize the intensity and duration of the enzyme treatment on a species-by-species basis along with other parameters in the respective protocols. Ultimately, the same reasoning should hold true – permeabilizing the cell wall will make it easier for biomolecules to pass through.

The surprising discovery of carrier DNA integrated into the termini of a once linear episome led us to explore the potential for assembly directly in *P. tricornutum*. It was possible to simultaneously electroporate multiple episomal fragments that could recombine through NHEJ or HR to form a single episome. The HR pathway is much more robust for assembly purposes, with ∼87% (26/30 colonies) screening positively for the recombined fragment junctions and 70% of recovered episomes having the expected sequence. This presents an exciting new opportunity for assembling *de novo* episomes directly in *P. tricornutum*, coined as **<u>d</u>**iatom **i**n **<u>v</u>**ivo **<u>a</u>**ssembly (DIVA), foregoing the need for traditional cloning steps in *E. coli* and *S. cerevisiae*; however, questions remain about the applications and limits of *in vivo* diatom assembly. There will be an upper limit to the number of fragments that can be simultaneously transformed, but we do not know what this limit is yet. We also do not know how large or small the fragments can be, and the ideal size range for the overlapping regions. The fragments used for HR assembly in this paper ranged from 6.8 to 3.0 kb and had ∼160 bp overlapping regions; in *S. cerevisiae*, constructs larger than 500 kb have been assembled from 25 overlapping ∼20 kb fragments before^31^, and successful recombination can be achieved with overlaps as small as 20 bp^32^. Going forward, it will be necessary to further characterize these parameters to make full use of DIVA.

Past methods for Cas9-based engineering in *P. tricornutum* included the delivery of RNPs through biolistics^23^ or the bacterial conjugation of episomes expressing the Cas9 complex^9^. Our paper adds electroporation and PEG transformation as additional tools for Cas9 genome engineering, with it being possible to deliver episomes through either method or directly electroporate RNPs into *P. tricornutum*. We generated thousands of 2-FA resistant transformants by electroporating an RNP complex targeting *PtAPT*, several-fold more efficient than what was previously reported with biolistics^23^. It was also possible to cotransform an episome and the *PtAPT* RNP complex at a rate of ∼10%. This offers an alternative method for engineering regions of the genome that are not counter-selectable, as a proportion of transformants that have uptaken the episome, which can be selected for, will have also uptaken the RNP and incurred genome editing. The capacity to perform multigene engineering by electroporating multiple RNP complexes was not explored in this study, but we believe this will be possible as this has been previously accomplished via biolistics^23^.

Complete sequencing data could not be captured for all the analyzed *PtAPT* mutants. This is likely due to the presence of large deletions that extend beyond the range of the primer binding locations. For reference, deletions as large as ∼2.7 kb have been observed in Cas9-edited *P. tricornutum* loci before^33^ and the primer sets for screening the *PtAPT* alleles were only positioned up to ∼450 bp away from the cut site. The alleles that could be sequenced demonstrated that Cas9 engineering had occurred in all assessed clones and revealed the unexpected insertion of carrier DNA in some clones. Based on this finding, we anticipate that it will be possible to intentionally introduce heterologous sequences into the *P. tricornutum* genome by simultaneous electroporation of an RNP along with a selectable DNA fragment bearing homology arms to the targeted locus. This could be used to create landing pads dispersed throughout the genome, facilitating multigene engineering at disparate loci, or to introduce yeast artificial chromosome (YAC) elements, allowing for the downstream capture of whole algal chromosomes via transformation-associated recombination (TAR) cloning (Fig. 7). Episome-based Cas9 genome engineering has been used to integrate heterologous sequences into the *P. tricornutum* genome before^34^, so we anticipate that RNP-based integration of similarly sized heterologous inserts should be possible.

**Figure 7.**
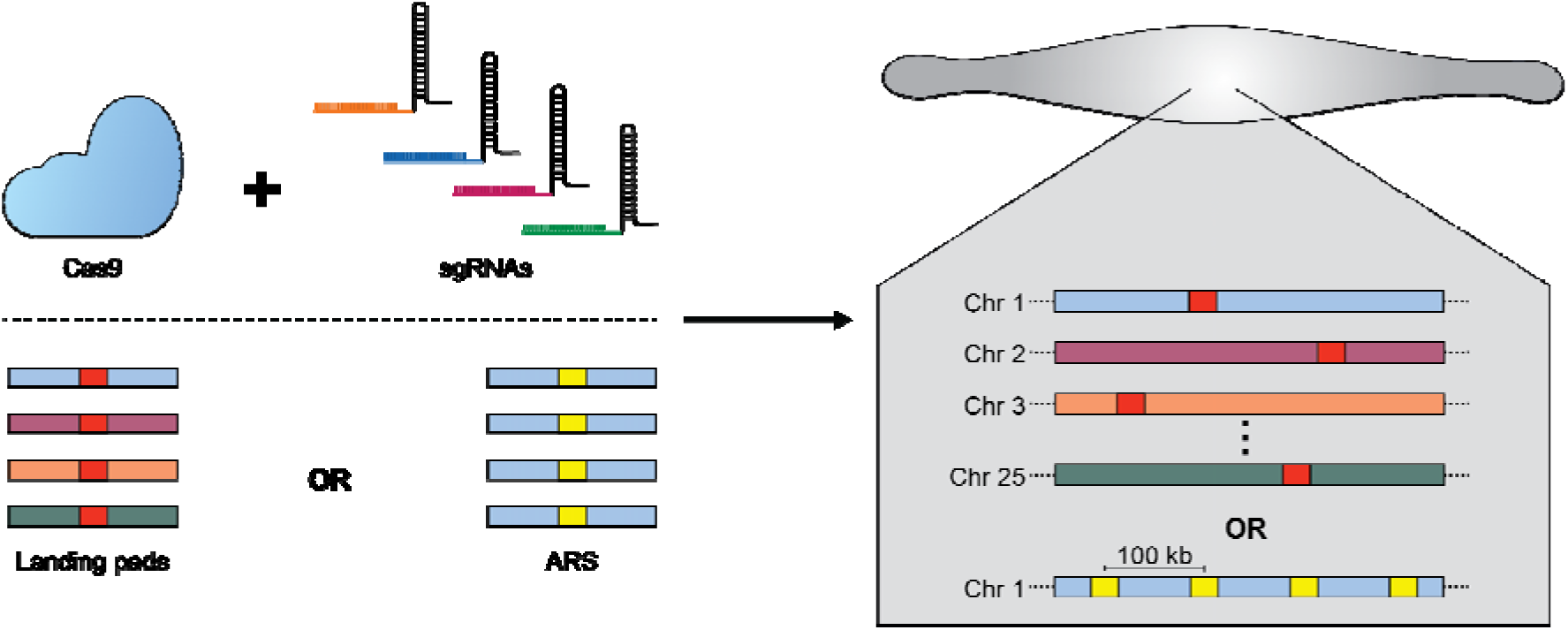
Installing landing pads or autonomous replication sequences (ARS) throughout the *P. tricornutum* genome. A Cas9 complex with multiple sgRNAs is simultaneously electroporated with donor template DNA fragments to introduce heterologous sequences into the *P. tricornutum* chromosome(s). Figure is adapted from and inspired by the work of Baek et al^35^.

The methods presented in this study pave the way for a new era of accelerated engineering in *P. tricornutum*. When we first proposed the Pt-syn 1.0 project, we envisioned using bacterial conjugation to introduce synthetic chromosomes into the algal cell^14^. Now, we can envision the *in vivo* assembly of synthetic chromosomes directly in the algal nucleus or the piece-by-piece replacement of endogenous chromosomes with synthetic parts (Fig. 8). With the potential to integrate YAC elements throughout the algal genome, we also foresee TAR cloning all nuclear chromosomes in *S. cerevisiae,* where they can be more readily tinkered with. Challenges and questions remain about the limits of these approaches that future work will have to address before these feats can be accomplished, but in any case, this work has led us one step closer towards actualizing Pt-syn 1.0 – a diatom controlled by a wholly-synthetic genome.

**Figure 8.**
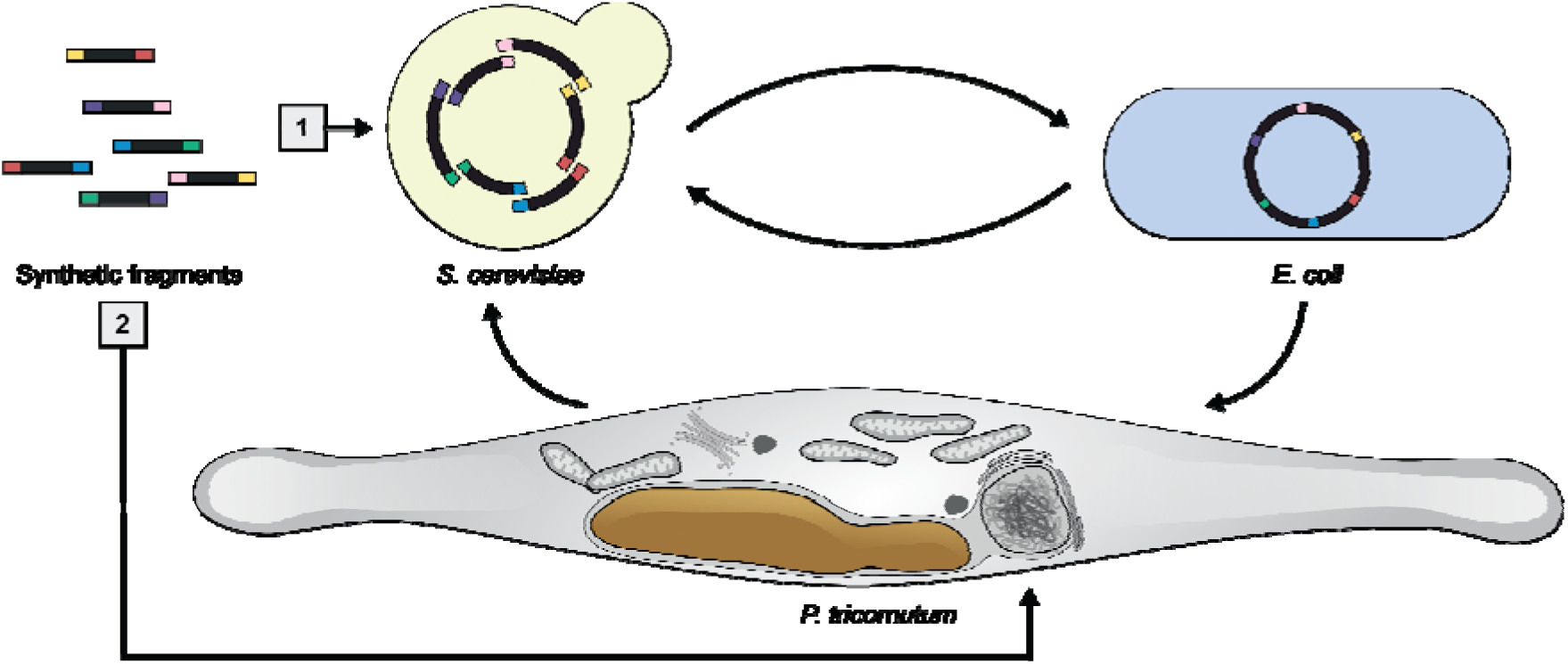
The canonical [1] and novel [2] pathways for delivering and assembling episomes in *P. tricornutum*. Prior to this work, overlapping synthetic fragments would be assembled in *S. cerevisiae*. Assembled episomes would be transferred to *E. coli* and then conjugated to *P. tricornutum*. In this pathway, the episome is shuffled between three host organisms during the design-build-test cycle. The novel pathway provides the opportunity to directly assemble episomes in the agal cell by electroporation of the synthetic fragments.

To conclude, this study presents optimized electroporation and PEG methods for high efficiency transformation of *P. tricornutum*, an emerging diatom chassis for algal synthetic biology. The techniques described in this manuscript have already been adapted for *T. pseudonana* and hold immense promise for other diatom species. The simplicity and scalability of these methods will be immensely valuable as the Pt-syn 1.0 project begins to build and test different synthetic chromosome designs.

## METHODS

### Microbial strains and growth conditions

1. *P. tricornutum* (Culture Collection of Algae, the University of Texas at Austin [UTEX], catalog number: 646) was cultured in liquid L1 media or on ½-salt L1 1% agar plates (w/v) in a growth chamber set to 18 °C and a light/dark cycle of 16/8 hours. Cultures were irradiated with white LED lights (Mars Hydro, model: TSL 2000) set to photon flux densities of ∼75 μEDm−2Ds−1 (liquid cultures) or ∼55 μEDm−2Ds−1 (agar plate cultures). Agar plates were placed in clear polystyrene trays to reduce desiccation. L1 media was prepared as previously described and lacks silica^6^.
2. *T. pseudonana* (Bigelow National Center for Marine Algae and Microbiota, catalog number: CCMP1335) was cultured in liquid F/2 media in a growth chamber set to 20 °C and 24 hour white light at a photon flux density of . Cultures were irradiation with X lights set to a photon flux density of 50 μEDm−2Ds−1.
3. *S. cerevisiae* strain VL6-48 (American Type Culture Collection [ATCC], catalog number: MYA-3666) was grown in 2X YPAD media or plated on complete minimal media lacking histidine with 1% agar (w/v) and 1 M D-sorbitol. Yeast cultures were incubated at 30 °C, with liquid cultures placed on a shaker set to 225 rpm. Agar plates were sealed with parafilm to reduce desiccation. Yeast media were prepared as previously described^36^.
4. *E. coli* strain EPI300 (LGC Biosearch Technologies, Lucigen, catalog number: EC300110) was grown in lysogeny broth (Miller formula) with or without 1% agar (w/v) and supplemented with 15 µg/ml chloramphenicol and/or 40 µg/ml gentamycin and/or 100 µg/ml L-arabinose. Cultures were placed in an incubator set to 37 °C, with liquid cultures placed on a shaker set to 225 rotations per minute (rpm).

### PCR amplification

Episomes and/or fragments for transformation and/or assembly were amplified with GXL polymerase (Takara, catalog number: R050A) using the rapid PCR protocol (all primers are listed in Supplementary Table S7). Of note, we used both phosphorylated and non-phosphorylated primers when amplifying pPtGE31_ΔPtR as a linear episome. PCR reactions were conducted with a total of 25 to 35 cycles depending on amplification efficiency. Fragments that were amplified from an episome template were treated with 10 units (0.5 μl) of *Dpn*I (New England Biolabs, catalog number: R0176) and incubated at 37 °C for 30 min before deactivation at 80°C for 20 min. After deactivation, fragments were column-purified using the PureLink PCR Purification Kit (Invitrogen, catalog number: K310001) and eluted with ddH_2_O.

### PCR screening of algal and E. coli transformants

DNA isolated from algal and *E. coli* transformants was screened for the presence of specific episomal regions using a PCR assay. Here, 1 μl of DNA isolated through alkaline lysis was used as template in a 10 μl SuperPlex PCR reaction (Takara, catalog number: 638543). For all algal transformants, episomes were analzyed with a single primer set at a time as there was rampant off-target binding when multiple primer pairs were used simultaneously (all primers are listed in Supplementary Table S7). We were able to successfully conduct multiplex PCR using *E. coli* template DNA; however, when screening the NHEJ assembled episomes, we chose to perform singleplex screens due to the unexpected recombination of the fragments. Reactions were carried out according to the manufacturer guidelines for a total of 30 cycles.

### DNA isolation

Episomes were isolated from *P. tricornutum*, *S. cerevisiae*, and *E. coli* using a modified alkaline lysis protocol that was previously described in Karas et al^6^. We modified steps 1 to 3 for *P. tricornutum* DNA isolation, which is described below.

<u>Steps 1 to 3 for *P. tricornutum*:</u> (**1**) *P. tricornutum* transformants that were repatched at least twice were grown to high density on a ¼-salt L1 plates supplemented with the appropriate antibiotic(s) (i.e., 100 µg/ml nourseothricin or zeocin for single selection, double selection plates contained both antibiotics at 50 µg/ml). Colonies were passaged as large streaks to ensure there were enough cells for DNA isolation – typically, a single plate would be used to passage 6 colonies that were struck out to each cover approximately a sixth of the plate. Plates were grown until a reasonable cell density was achieved, which typically took 4 days for nourseothricin plates and upwards of 6 days for zeocin or double-selection plates. (**2**) Cells were scraped and resuspend in 250 µl of resuspension buffer, which consisted of 245 µl of P1 (Qiagen, catalog number: 19051) and 5 µl of alcalase (Sigma-Aldrich, catalog number: 126741). (**3**) The resuspended cell mixture was vortexed for 1 to 3 seconds, then placed in a heating block set to 56 °C for 10 minutes – this is within the optimal temperature range for alcalase activity, according to the manufacturer. After 10 minutes passed, the cells were vortexed again for 1 to 3 seconds to ensure there were no cell clumps.

For *E. coli*-derived episomes smaller than 11 kb, we used column purification kits to isolate and purify DNA (New England Biolabs, catalog number: T1110). DNA was eluted with ddH_2_O equilibrated to 56 °C.

To isolate a high concentration of pSC5 (∼56 kb) from *E. coli*, alkaline lysis was performed using 27 ml of culture (1.5 ml culture per lysis reaction, 18 reactions in total). The DNA pellets were resuspended with ∼25 µl of sddH_2_O and combined into a single 1.5 ml tube (∼460 µl total). Half of the isolated DNA was then digested using *Sac*I-HF, as described below. Then, a tenth volume of sodium acetate (3M, pH 5.2) and two volumes of ice-cold 100% ethanol were added to the samples. The solutions were inverted to mix and placed into a -80 °C freezer for an hour before performing centrifugation at 16,000 x G for 10 minutes at 4 °C. The DNA pellets were then washed twice with 500 µl of ice-cold 70% ethanol before being decanted, dried, and resuspended with 50 µl of sddH_2_O. This additional sodium precipitation step was performed to further concentrate the samples and reduce the carryover of contaminants, particularly from the digest reaction.

DNA concentrations were measured by fluorometry using the DeNovix dsDNA Broad Range Assay (FroggaBio, catalog number: DSDNA-BROAD-EVAL) and purity ratios were assessed using a Nanodrop spectrophotometer. For the experiments shown in Fig. 2 to 4, samples were adjusted to be equimolar prior to conducting electroporation.

### Restriction digests of episomes

Circular pPtGE31_ ΔPtR was isolated a *P. tricornutum* transformant and recovered in EPI300 *E. coli*. After isolating the episome from *E. coli*, approximately 10 µg was digested using 3 µl of *Eco*RI-HF (New England Biolabs, catalog number: R3101) at 37 °C for 1 hour. An additional reaction was conducted where 10 µg was digested using 3 µl of *Eco*RI-HF and 3 µl of mung bean nuclease (New England Biolabs, catalog number: M0250) at 37 °C for 1 hour. Complete digestion was confirmed by agarose gel analysis, then samples were purified using a DNA Cleanup Kit (New England Biolabs, catalog number: T1030).

Circular pSC5 was isolated from EPI300 *E. coli* as described above. Prior to conducting the sodium acetate precipitation step, approximately 40 µg of pSC5 was digested with 18 µl of *Sac*I-HF (New England Biolabs, R3156) at 37 °C for 1 hour. Complete digestion was confirmed by agarose gel analysis, then the sample was purified by sodium acetate precipitation, as described above.

### Preparation of Cas9:sgAP1 complex

HiFi Cas9 nuclease (catalog number: 1081060) and synthetic sgRNA were purchased from Integrated DNA Technologies (IDT) and prepared according to the manufacturer’s guidelines. Briefly, 2 to 4 µg of Cas9 (62 µM) was combined with sgRNA (100 µM) in a 1:1.2 molar ratio and then incubated at room temperature for 20 minutes to form the Cas9:sgRNA complex. After incubation, the complex was set on ice for up to 1 hour, during which the *P. tricornutum* cells were prepared for electroporation. Once the cells were ready, 40 µg of ssssDNA was added to a 50 µl preparation of the cells, followed by the Cas9:sgRNA complex. The mixture was pipetted up-and-down a few times before transferring to an electrocuvette and promptly pulsed. For the co-transformation experiment, we added 500 ng of PCR-amplified pPtGE31_ ΔPtR to the cells ahead of electroporation.

### Yeast assembly of pPtGE31_ShBle

The episome pPtGE31_ShBle was created by yeast assembly of three overlapping fragments. First, the episome pPtGE31_ ΔPtR was amplified as two fragments using primers that would add 40 bp overlaps for the promoter and terminator region of the ShBle cassette and split the episome in the HIS3 region. Then, the ShBle cassette was amplified from the p0521s plasmid (Addgene ID: 62862)^6^ using primers that would add 40 bp overlaps for the insertion site in pPtGE31_ ΔPtR (all primers are listed in Supplementary Table S7). Yeast assembly was conducted as previously described^36^ using ≥ 100 ng of each fragment in place of a bacterial donor.

### Electroporation to P. tricornutum

*P. tricornutum* cells were grown in liquid media or on agar plates for the experiments conducted in Fig. 1 – all successive experiments used cells derived from agar plates. For liquid cultures, 1 ml of cells adjusted to 1 – 2 x 10^8^ cells/ml were passaged into 50 ml of L1 media and placed in a growth chamber on a shaker set to ∼100 rpm. For agar plate cultures, 250 µl of cells adjusted to 1 x 10^8^ cells were spread onto a ½-salt L1 plate with 1% agar (w/v). Liquid and agar plate cultures were grown for 4 days. After 4 days, cells were scraped from the agar plates using 750 µl of L1 and transferred to a 1.5 ml tube. An additional 750 µl of L1 was used to remove any remaining cells from the plate. Then, 10^-1^ to 10^-2^ dilutions of the liquid and agar plate cultures were prepared and used to estimate cell density with a hemocytomer. Liquid and agar plate cultures were adjusted to have a final cell density between 2 – 4 x 10^8^ cells/ml; for the liquid cultures, this requires concentrating the cells via centrifugation and resuspension in a smaller volume. Here, the cells are transferred from the flask to a 50 ml conical tube and pelleted by centrifugation at 2000 x G for 10 minutes at 18 °C. The pellet was resuspended with 375 mM D-sorbitol to a concentration between 2 – 4 x 10^8^ cells/ml and transferred to a 1.5 ml tube. Cells derived from agar plates were pelleted by centrifugation at 750 x G for 4 minutes; here, we used a microcentrifuge, which could pellet the cells at a lower centrifugal speed. As before, the cells were resuspended using 375 mM D-sorbitol to a concentration between 2 – 4 x 10^8^ cells/ml. The final cell concentration was equalized between the liquid and agar plate derived cultures for every biological replicate.

To spheroplast, 1 mAnson of alcalase (Sigma-Aldrich, catalog number: 126741) was added for every 1 x 10^8^ cells. At this stage, the cells will have been previously concentrated to a density between 2 – 4 x 10^8^ cells/ml and already resuspended in 375 mM D-sorbitol. Alcalase was diluted 10-times in sddH_2_O to reduce the pipetting error associated with small volumes. After the addition of alcalase, cultures were placed on a rocking shaker set to ∼15 rpm at room temperature for 20 minutes, during which spheroplasting occurs. Once 20 minutes has passed, the cells were pelleted by centrifugation at 325 x G for 4 minutes. The supernatant is decanted, and the cells are gently resuspended with 1 ml of ice-cold 375 mM D-sorbitol using a P1000 pipette.

The electroporation protocol we used is adapted from Zhang and Hu^5^, with adjustments made by Kassaw et al^16^ and Pampuch (unpublished). All materials (i.e., cells, DNA, D-sorbitol, electrocuvettes) should be kept on ice for the duration of this protocol. First, the spheroplasted cells were pelleted by centrifugation at 325 x G for 4 minutes. The supernatant is decanted, and the pellet is gently resuspended with 1 ml of ice-cold 375 mM D-sorbitol. This process of centrifugation and resuspension is repeated another 2 times, such that in total, the cells will have been resuspended with 375 mM D-sorbitol five times (this includes centrifugation steps prior to and post spheroplasting). After the fifth wash, the cells are spun at 750 x G for 3 minutes and a P200 pipette is used to gently remove any remaining supernatant. The cells are gently resuspended with 100 to 200 µl of 375 mM D-sorbitol such that the total volume of resuspended cells will be between ∼150 to 250 µl. Into sterile 1.5 ml tubes, 50 µl of cells are aliquoted along with 4 µl (10 µg/µl) of single stranded salmon sperm DNA (ssssDNA, Sigma-Aldrich, catalog number: D7656) and the DNA to be transferred. Of note, the ssssDNA should be heated to 95 °C for 2 to 5 minutes and then cooled on ice prior to this to ensure it remains single stranded. We used between 1 to 1000 ng of DNA for all transformations. Negative controls consisted of only ssssDNA in the cell mixture.

Cells are gently pipetted up and down 2 times before transferring to an ice-cold sterile electrocuvette (Fisher Scientific, catalog number: FB102), ensuring that no air bubbles have been transferred and the cells lie evenly across the bottom. The electrocuvette is pulsed in an electroporator (BioRad, Gene Pulser Xcell System, catalog number: 1652660) set to the following parameters – voltage: 500 V, capacitance: 50 μF, and resistance: 400 Ω. The time constant varied from 30 to 32.5 milliseconds for the experiments conducted in Figures 1 to 4. Immediately after pulsing, 1 ml of L1 media is gently added to the electrocuvette without disturbing the cells. The electrocuvettes are left to rest at room temperature for 10 minutes before fully resuspending the cells and transferring the mixture to 10 ml of L1 in 50 ml falcon tubes. The lids of the falcon tubes are left loose for air transfer, and the cultures are moved to a growth chamber to recover for 16 to 24 hours.

After recovery, the cultures are pelleted by centrifugation at 2000 x G for 10 minutes at 18 °C. The pellet is resuspended with 500 µl of L1 media, then 50 µl (one-tenth), 125 µl (a quarter), and/or 250 µl (half) of the cells are spread onto ¼-salt L1 plates supplemented with the appropriate antibiotic(s).

It is worth noting that there is a great degree of variance between electroporation equipment, and this can drastically impact the efficiency of transformation to *P. tricornutum*. If others attempt this protocol, it may be necessary to initially attempt electroporation with several different parameters to determine which settings work best with different set-ups. For our set-up (500V, 50 μF, 400 Ω), an empty electrocuvette generates a time constant of ∼35 milliseconds. A neighbouring lab has successfully performed electroporation to *P. tricornutum* using an older electroporator set to the originally described parameters (500V, 25 μF, 400 Ω), which generates a time constant of ∼20 ms with an empty cuvette.

### PEG transformation of P. tricornutum

The optimized PEG transformation protocol is similar to the methods3f described by Karas et al.^6^, with some important modifications. Firstly, 2-4 x 10^8^ cells were protoplasted using 1000 µl of alcalase in place of the zymolyase, hemicellulase, and lysozyme cocktail. The cells and protoplasting solution (9 ml L1, 1 ml of alcalase) were combined in a 50 ml conical tube that was placed on a slow rocking shaker (∼15 rpm) and incubated at room temperature for an hour. After an hour, 40 ml of L1 was gently added to the cells which were then pelleted by centrifugation at 1500 x G for 5 minutes at 18°C. The supernatant was carefully decanted and the cells were resuspended with 500 µl of L1 followed by 8 µl of ssssDNA (10 µg/µl). Then, 250 µl of protoplasts were combined with episomal DNA (circular or linear, 500 ng to 1 μg) and 1 ml of 20% PEG-8000 (w/v; also contains 10 mM Tris pH 8, 10DmM CaCl_2_, 2.5DmM MgCl_2_) equilibrated to 30°C. This was gently mixed through inverting and then incubated at RT for 30 minutes. After incubation, the mixture was centrifuged for 7 min at 1500 x G. The supernatant was removed and cells were resuspended in 10 ml of L1 and recovered in the growth chamber for up to 48 hours. After recovery, the cells were centrifuged for 5Dminutes at 1500 x G and 18°C. Then the cells were resuspended in 500 μl of L1 and 125-250 μl was plated on ¼ × L1 supplemented with nourseothricin (100 μg/ml).

### PEG transformation of T. pseudonana

Approximately 1 x 10^5^ cells/ml of *T. pseudonana* was inoculated into f/2 medium. After 4 days of growth (end of exponential phase), cells were pelleted by centrifugation at 1500 x G for 10 minutes and resuspended to a concentration of 3 x 10^8^ cells/ml. The same transformation protocol was followed as described for *P. tricornutum*; however, the protoplasting step was conducted in f/2 medium with differing amounts of alcalase and incubation lengths (Fig. S12A). Approximately 500 ng of the 8.2 kb episome pBIG1 was combined with 40 µg of carrier DNA for transformation. Following PEG transformation, the cells were recovered in 10 ml of f/2 media and recovered for 2 days prior to plating. Post-recovery, the cells were centrifuged for 5 minutes at 1500 x G. The cells were then resuspended in 500 μl of f/2 media and 250 μl was plated on full-salt f/2 agarose plates supplemented with nourseothricin (100 μg/ml).

### Screening P. tricornutum colonies transformed with two fragments – NHEJ pathway

Non-overlapping fragments from pPtGE31_ShBle were electroporated both individually and simultaneously into *P. tricornutum* (Supplementary Fig. S3). Cells that were electroporated with both fragments were plated onto ¼-salt L1 plates supplemented with 100 µg/ml nourseothricin or 100 µg/ml zeocin. Transformants that appeared following electroporation with both fragments were passaged twice – first, onto media supplemented with the same antibiotic as the initial transformation, and then onto media supplemented with the alternative antibiotic (Supplementary Table S5). Twenty transformants, ten derived from each initial transformation plate, that demonstrated resistance to both nourseothricin and zeocin were passaged for a third time onto double-selection media (¼-salt L1 supplemented with 50 µg/ml nourseothricin and zeocin). DNA isolation was performed using cells from this third passage, followed by PCR screening.

### Screening P. tricornutum colonies transformed with multiple fragments – HR pathway

Overlapping fragments were adjusted to a final concentration of 56 nM, which is four-times more concentrated than the linear episome that was used as a positive control in these experiments (14 nM). Different combinations of the fragments were electroporated to measure the amount of carry-over for the template plasmid, pPtGE31_ShBle. Cells were plated onto ¼-salt L1 plates supplemented with 100 µg/ml nourseothricin and grown for at least two weeks before passaging 10 colonies from the assembly plates onto the same type of selection plate. The repatched colonies were grown for one week before passaging again for DNA isolation.

### Recovery of P. tricornutum episomes in E. coli

Episomes were isolated from *P. tricornutum* transformants as described above. For each electroporation reaction, 2 to 4 μl of *P. tricornutum* DNA was combined with 25 to 50 μl of homemade electrocompetent EPI300 cells (derived from LGC Biosearch Technologies, catalog number: EC02T110) in a sterile 1.5 ml tube. The mixture was gently pipetted twice and then transferred to a sterile 2 mm electrocuvette, ensuring no air bubbles were transferred. The electrocuvette is pulsed in an electroporator set to the following parameters – voltage: 2.5 kV, capacitance: 25 μF, and resistance: 200 Ω. This generated a time constant of 4.9 to 5.2 milliseconds. Electroporated cells were resuspended by pipetting 1000 μl of SOC media up-and-down in the electrocuvette, and then transferred to sterile 1.5 ml tubes to recover at 37 °C and 225 rpm for approximately 1 hour. After recovery, half of the reaction volume (∼400 μl) was spread across a 1.5% agar (w/v) LB plate supplemented with 15 μg/ml chloramphenicol (pPtGE31_ ΔPtR and pPtGE31_ShBle recovery) or 40 μg/ml gentamycin (pSC5 recovery), which were then transferred to a 37 °C incubator. Transformants appeared across the plates within 24 hours, with there being a range from 4 to ∼100 colonies per transformation.

<u>For recovery of pPtGE31_ ΔPtR and pPtGE31_ShBle episomes</u>: single *E. coli* colonies were picked and inoculated into 5 ml of LB media supplemented with 15 μg/ml chloramphenicol. Cultures were grown overnight at 37 °C and 225 rpm. The following day, 500 μl of culture was inoculated into 4.5 ml of LB media supplemented with 15 μg/ml chloramphenicol and 100 μg/ml L-arabinose. These episomes contain a pCC1BAC backbone, which can be induced to high-copy number replication in EPI300 through by adding L-arabinose to the media. Induced cultures were grown for at least five hours prior to episome isolation, with 1.5 ml of culture being used for DNA isolation, as described above.

<u>For recovery of the conjugative plasmid pSC5</u>: DNA isolated from eight *P. tricornutum* pSC5 transformants were electroporated into EPI300 cells. Four colonies per transformation were inoculated into 5 ml of LB supplemented with 40 μg/ml gentamycin. Additionally, EPI300 strains carrying pSC5 (positive control) or pSAP (recipient cell line) were inoculated from glycerol stocks into 5 ml of LB supplemented with 40 μg/ml gentamycin (pSC5) or 15 μg/ml chloramphenicol (pSAP). The cultures were placed at 37 °C and 225 rpm. After the cultures reached saturation (∼18 hours), 125 μl of the pooled *E. coli* transformants and pSC5 cultures were inoculated into 5 ml of LB supplemented with 40 μg/ml gentamycin. For the pSAP culture, 5 ml of culture was diluted into 45 ml of LB supplemented with 15 μg/ml chloramphenicol. The cultures were grown at 37 °C and 225 rpm for 2 hours. Then, the cultures were pelleted by centrifugation at 3000 x G for 12 minutes at 10 °C, decanting the supernatant post centrifugation. The pooled transformants and pSC5 cultures were each resuspended with 100 μl of 10% glycerol, whereas the 50 ml pSAP culture was resuspended with 1000 μl of glycerol. Strains were kept on ice during these steps to cease bacterial growth.

For conjugation, 100 μl of the donor cells (transformant pools [x8] and pSC5) were mixed with 100 μl of the recipient cells (pSAP). The cells were mixed by pipetting up and down 4 times before transferring the whole mixture to a 2% agar (w/v) LB plate. Additionally, 100 μl of the recipient was plated by itself to serve as a negative control. Conjugation plates were incubated at 37 °C for 1 hour. Then, plates were scraped using 2 ml of sddH_2_O. The scraped cells were vortexed for 5 seconds to ensure any clumps were dispersed. Transconjugants were spot plated onto 1.5% agar (w/v) LB plates supplemented with 40 μg/ml gentamycin and 15 μg/ml chloramphenicol with dilutions ranging from 10^0^ to 10^-8^.

### Sequencing and alignment

Amplicons generated by amplifying the episome termini region in *P. tricornutum* transformants were sent for Sanger sequencing (London Regional Genomics Centre). Episomes isolated from *E. coli* were sent for Oxford Nanopore whole-plasmid sequencing (Plasmidsaurus Inc). The resulting sequences were manually aligned to the respective template episomes. Regions that could not be manually aligned were queried using a local alignment algorithm (EMBOSS water tool)^37^ to search for homology in the episome. Sequences that could not be aligned manually or algorithmically were queried using BLASTn (NCBI)^38^.

### Analyses of P. tricornutum transformation efficiency

The total number of colonies per transformation reaction were estimated by manually counting the number of CFUs per plate and then multiplying this value by 2 or 10, depending on whether half or a tenth of the total reaction was plated, respectively. In instances where the transformation plate was too dense to reliably count by eye, five 1 cm^2^ quadrants were sampled from the plate and counted using a dissecting microscope with 10x magnification. Here, the total number of CFUs was estimated by averaging the number of colonies per quadrant and multiplying this value (i.e., colonies per cm^2^) by the total surface area of the petri dish.

Transformation efficiencies were estimated by dividing the total number of CFUs by the number of cells per reaction. When graphically comparing transformation reactions, all calculations and plotting were done in R; otherwise, we used an Excel spreadsheet keep track of data and tabulate transformation efficiencies. The values for the standard error of the mean were calculated by dividing the standard deviation for a series of experiments by the square root of the number of biological replicates.

## ACKNOWLEDGEMENTS

This work was supported by Natural Sciences and Engineering Research Council of Canada (RGPIN-2018-06172 and RGPIN-2025-05428) awarded to B.J.K. and funding from the Natural Environment Research Council (NERC) grant NE/Z504130/1 awarded to T.M. We would like to acknowledge Drs Graham Peers and Tessema Kassaw for sharing their knowledge on electroporation to *P. tricornutum* with us. Our explorations into this method began with a protocol that was shared by Dr. Kassaw, who assisted us with initial troubleshooting. We would also like to acknowledge Karen Nygard, a microscopy specialist at Western University’s Biotron Facilities, who assisted with capturing the microscopic images shown in Fig. 1A and B.

## AUTHOR CONTRIBUTIONS

MP and GT established the parameters necessary for conducting electroporation in the Karas lab and performed experiments 1 through 79 in Supplementary Table 1. EJW established the spheroplasting technique for electroporation and PEG transformation in *P. tricornutum*. She also performed all other *P. tricornutum* experiments detailed in the manuscript. LD and YL performed the PEG transformation experiments in *T. pseudonana* and generated the images used for Fig. S12. EJW wrote the manuscript and created all other figures in Adobe Illustrator; MP, LD, YL, TM, and BJK edited the manuscript and provided crucial insight. BJK supervised all experimentation done by EJW, MP, and GT, and assisted with experimental design, data analysis, troubleshooting, and more.

## COMPETING INTEREST DECLARATION

The authors declare no competing interests.

## Notes

### Competing Interest Statement

The authors have declared no competing interest.

### Summary of Updates

We have added multiple additional results, as well as three authors. New results: 1)The first demonstration that episomes can be assembled through homologous recombination in the diatom cell. 2)High-efficiency electroporation of a Cas9 ribonucleoprotein complex into the diatom nucleus for DNA-free engineering. 3)Co-transformation of episomes with Cas9 ribonucleoprotein complexes. 4)Establishment of a high-effeciency PEG transformation method for P. tricornutum. 5)Adaptation of the PEG transformation method to Thalassiosira pseudonana, another industrially relevant diatom species.

