## Supplemental file for "Spheroplasted cells: a game changer for DNA delivery to diatoms"

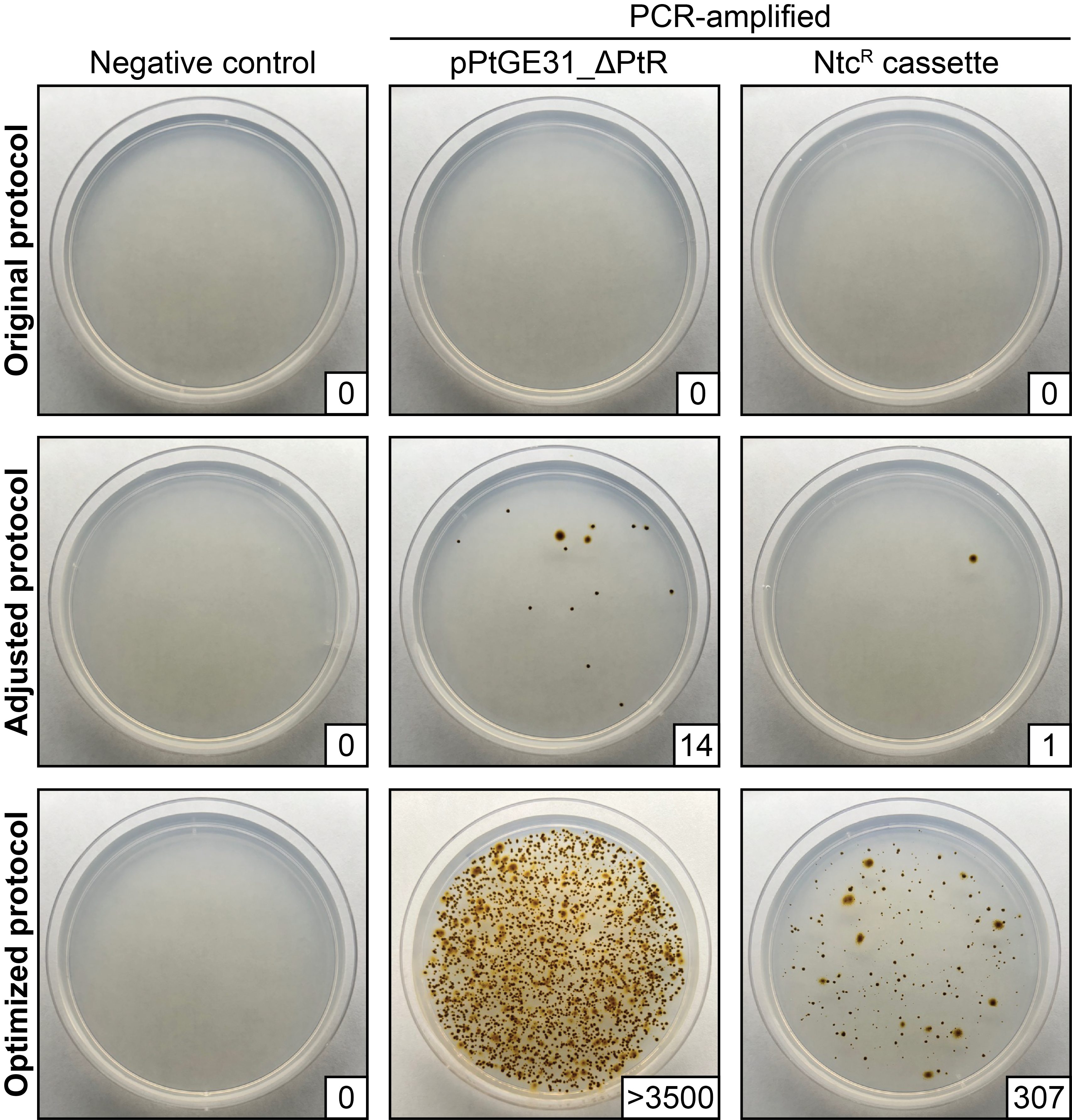

**Figure S1.** A comparison between the electroporation protocols described in the main text. We began our explorations with a protocol described by Kassaw et al^1^ (adapted from the methods of Zhang and Hu^2^; referred to as the original protocol). This protocol did not work in our experimental set-up until the capacitance parameter of the electroporator was adjusted from 25 µF to 50 µF (referred to as adjusted protocol). Adding an additional spheroplasting step ahead of electroporation drastically increases the efficiency for both episomal and integrative DNA (referred to as the optimized protocol). Experiments were conducted on the same day using the same number of cells (2 x 10^8^ per reaction) and concentrations of DNA. The Ntc^R^ cassette consists of the nourseothricin resistance gene flanked by the algal FcpD/FcpA promoter/terminator pair. A quarter of the reaction was plated on ¼-salt L1 plates supplemented with 100 µg/ml nourseothricin (NTC). Equimolar amounts of episome (1 µg, 11 kb) and the cassette (~130 ng, 1.4 kb) were used for all the reactions.

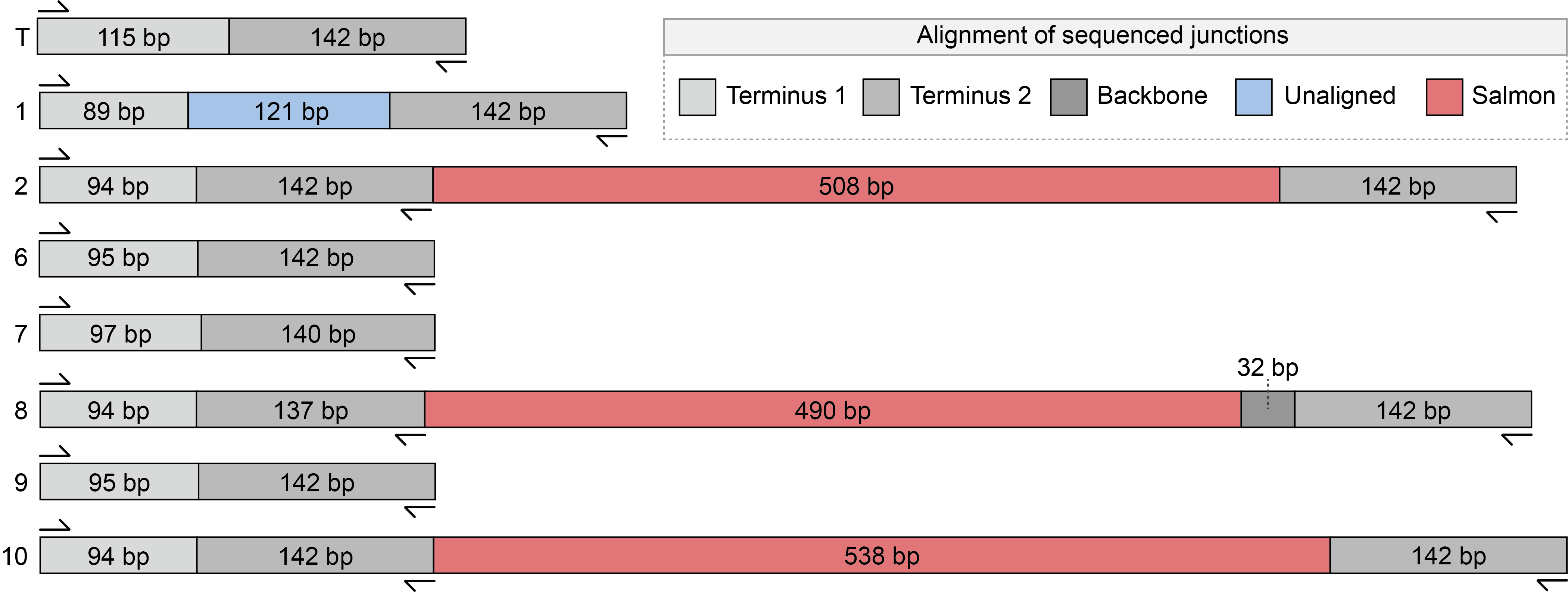

**Figure S2.** Sanger sequencing of the junction region for *P. tricornutum* cells transformed with PCR-amplified pPtGE31_ ΔPtR. Sequences were aligned to the template (T), which contains the region where the episome was split during PCR-amplification. The sequencing primers were positioned ≥100 bp away from the termini to capture if any insertions or deletions were occurring in this region. Colonies 1, 2, 8, and 10 demonstrate insertions of 121 to 538 bp in size. The insertion in colony 1 did not demonstrate complementarity to any regions of pPtGE31_ ΔPtR, nor did it have any BLASTn results. The 490 to 538 bp insertions in colonies 2, 8, and 10 were queried with BLASTn and demonstrated sequence similarity (≥88%) to various regions of the salmon and/or trout genomes. Deletions of 2 to 47 bp in length occurred in colonies 6, 7, and 9.

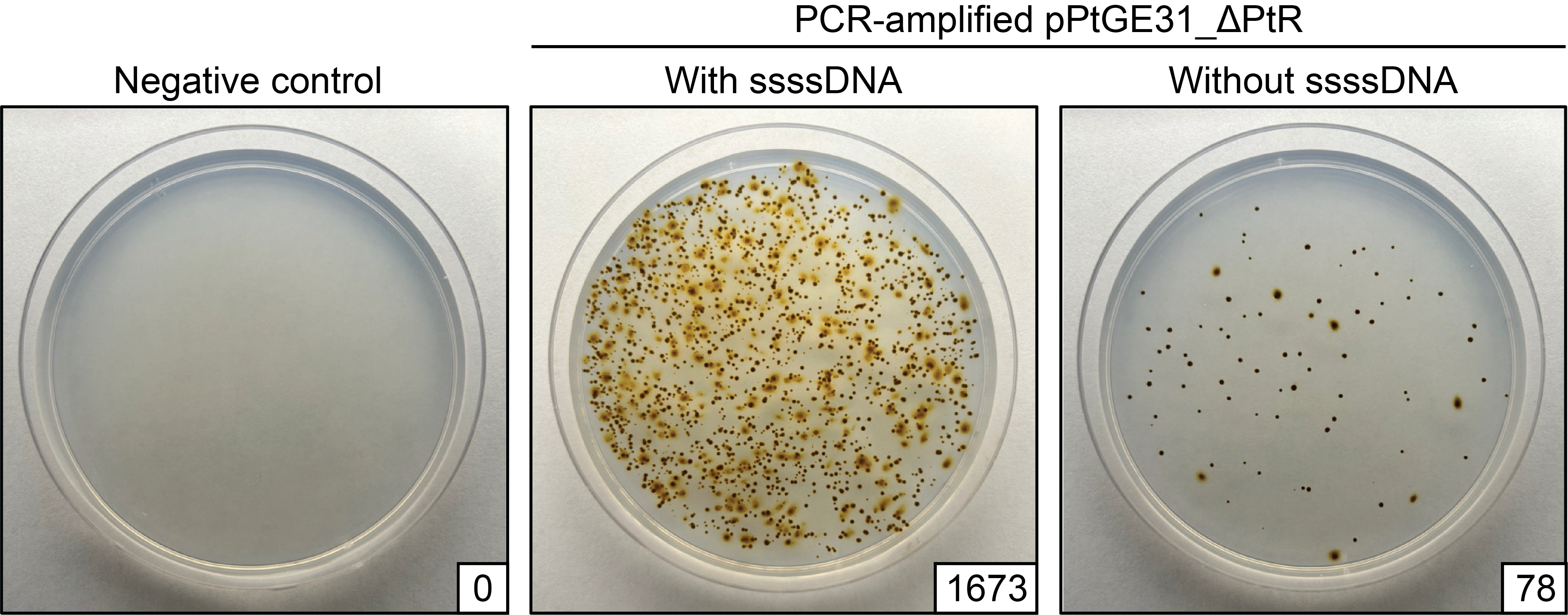

**Figure S3.** Electroporation of PCR-amplified pPtGE31_ ΔPtR with and without the addition of 40 µg of single stranded salmon sperm (ssss) DNA per reaction. The number of colony forming units (CFUs) is depicted in the bottom right corner for each transformation plate. Following electroporation, one-tenth of the total reaction was plated on ¼-salt L1 plates supplemented with 100 µg/ml nourseothricin. When this experiment was repeated with three biological replicates, the difference in efficiency was approximately 15-fold, on average.

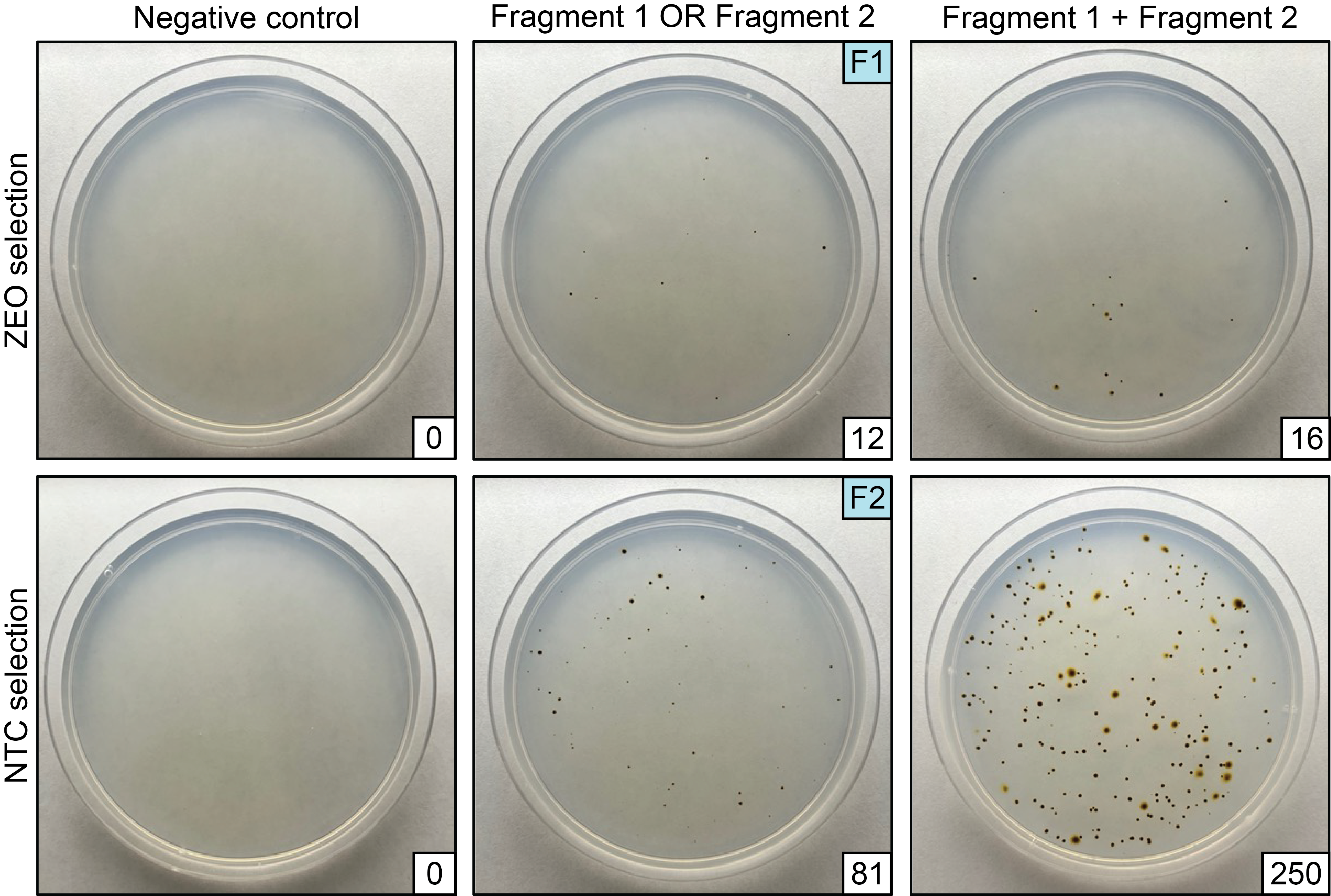

**Figure S4.** Electroporation of two fragments into *P. tricornutum*. Fragment 1 contains a zeocin resistance marker, whereas fragment 2 contains a nourseothricin resistance marker. Electroporation was conducted using the individual fragments and both fragments at once. The number of colony forming units (CFUs) is depicted in the bottom right corner for each transformation plate. For each electroporation, half of the total reaction was plated on ¼-salt L1 plates supplemented with 100 µg/ml nourseothricin (NTC) or zeocin (ZEO).

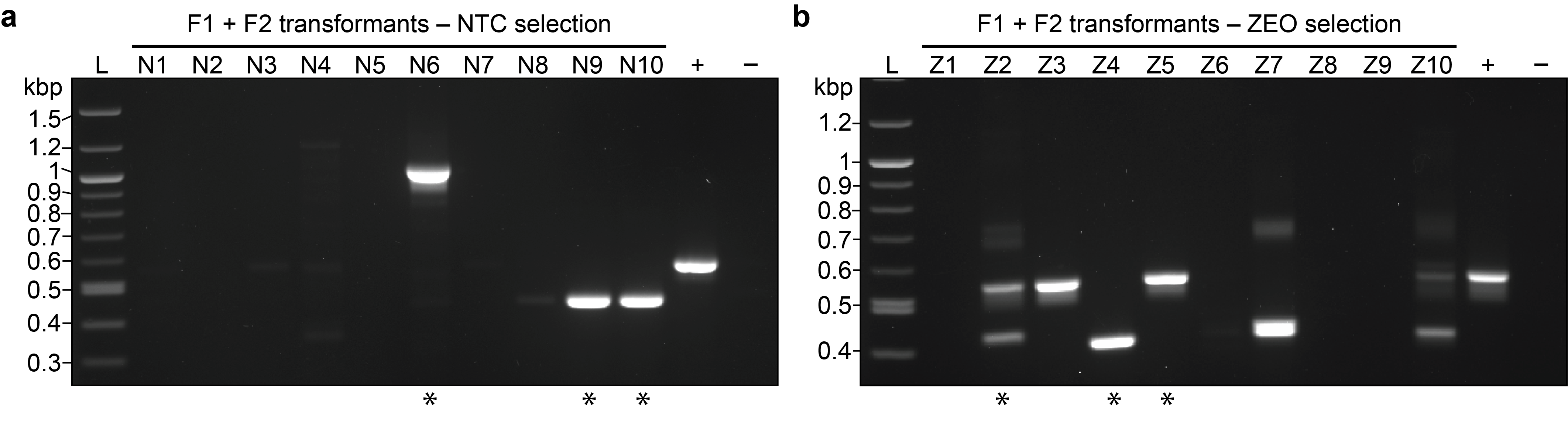

**Figure S5.** Screening *P. tricornutum* transformants following electroporation with two non-overlapping fragments simultaneously. (**a**) A PCR screen for junction B across ten algal transformants that were originally passaged from the transformation plate supplemented with 100 µg/ml nourseothricin (NTC) or (**b**) 100 µg/ml zeocin (ZEO). Dilute pPtGE31_ ShBle and genomic *P. tricornutum* DNA were used for the positive and negative controls, respectively. Single asterisks (*) indicate the algal colonies that had their DNA isolated and transformed into *E. coli*.

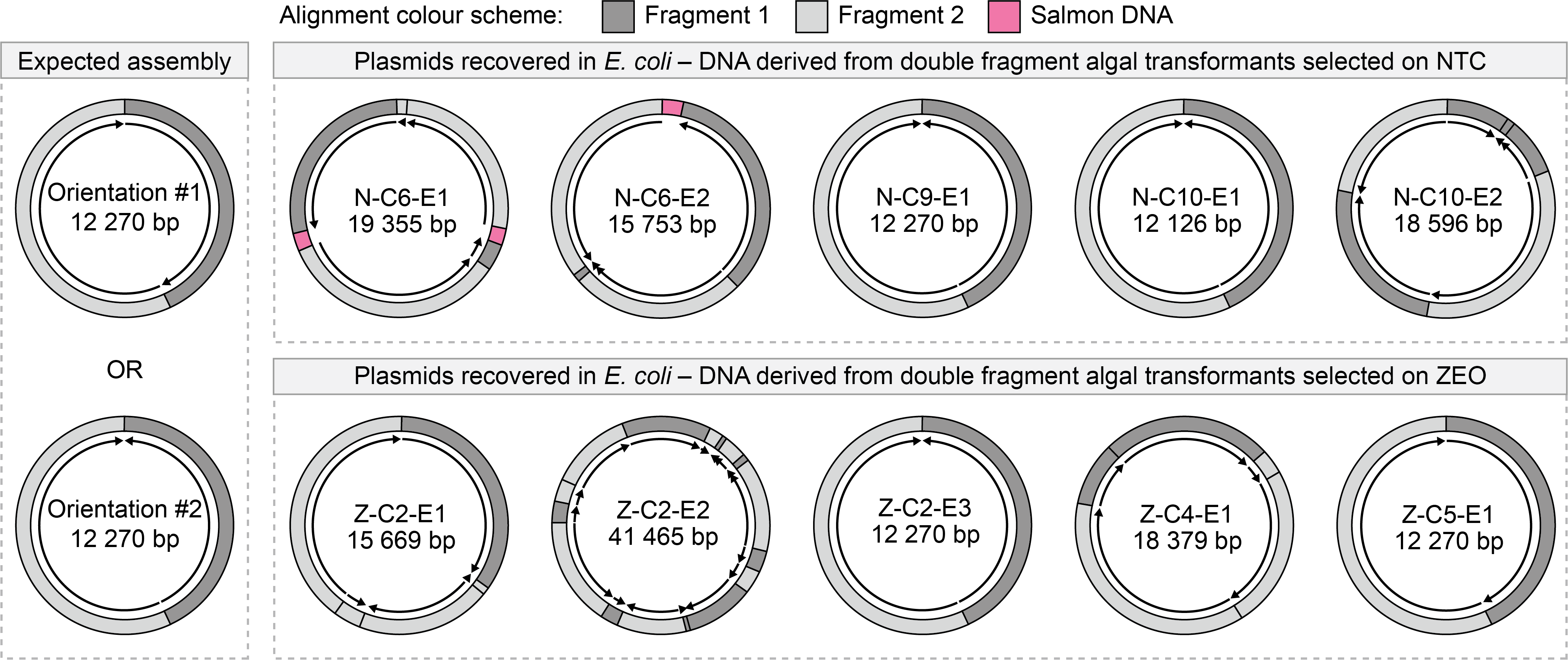

**Figure S6.** Sequencing of *P. tricornutum* NHEJ-assembled episomes recovered from *E. coli* transformants. Fragment orientations are depicted by arrows and are relative to the DNA directionality in the original plasmid, pPtGE31_ShBle.

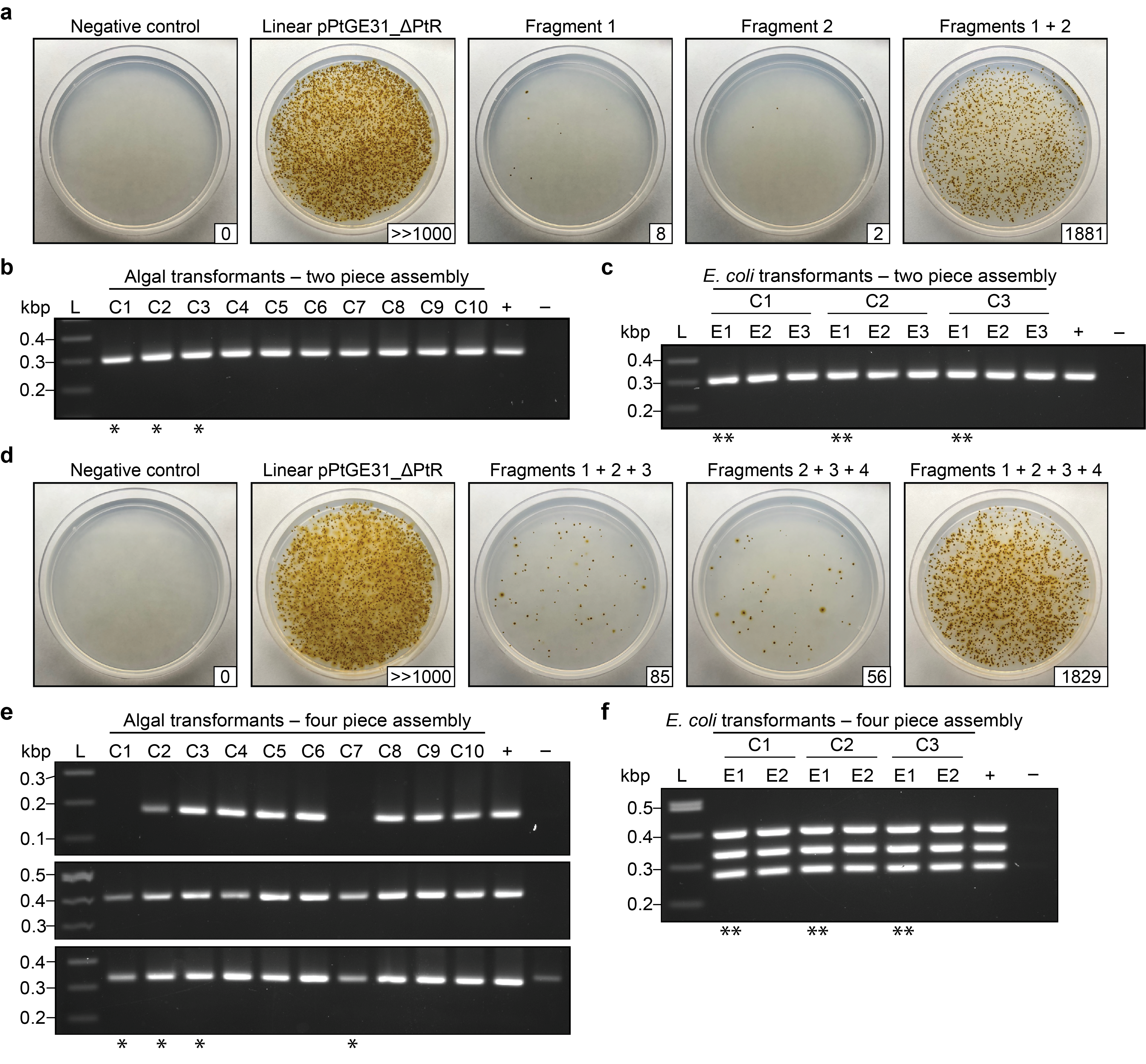

**Figure S7.** Electroporation and assembly of overlapping fragments in *P. tricornutum*. (**a**) Transformation of two overlapping fragments via electroporation. The fragments were also electroporated individually to determine the rate of carry-over of the template episome. (**b**) Screening 10 algal transformants post electroporation with two overlapping fragments and (**c**) nine *E. coli* transformants with episomes recovered from C1, C2, and C3. (**d**) Transformation of four overlapping fragments via electroporation. The fragments were also electroporated in combinations (i.e., F1 + F2 + F3 or F2 + F3 + F4) to determine the rate of carry-over of the template episome. (**e**) Screening 10 algal transformants post electroporation with four overlapping fragments and (**f**) six *E. coli* transformants with episomes recovered from C1, C2, and C3. For all transformations, linear pPtGE31_ ΔPtR (500 ng) was used as a positive control and half of the total reaction was plated on ¼-salt L1 plates supplemented with 100 µg/ml nourseothricin. The number of CFUs is shown in the bottom right corner for each plate. For algal screens, the positive control consists of dilute pPtGE31_ShBle, and the negative control consists of wild-type *P. tricornutum* DNA. The same positive control was used for E. coli screens, however, the negative control consisted of double-distilled water. Single asterisks (*) indicate the algal colonies that had their DNA isolated and transformed into *E. coli*. Double asterisks (**) indicate the *E. coli* colonies that were sent for whole plasmid sequencing.

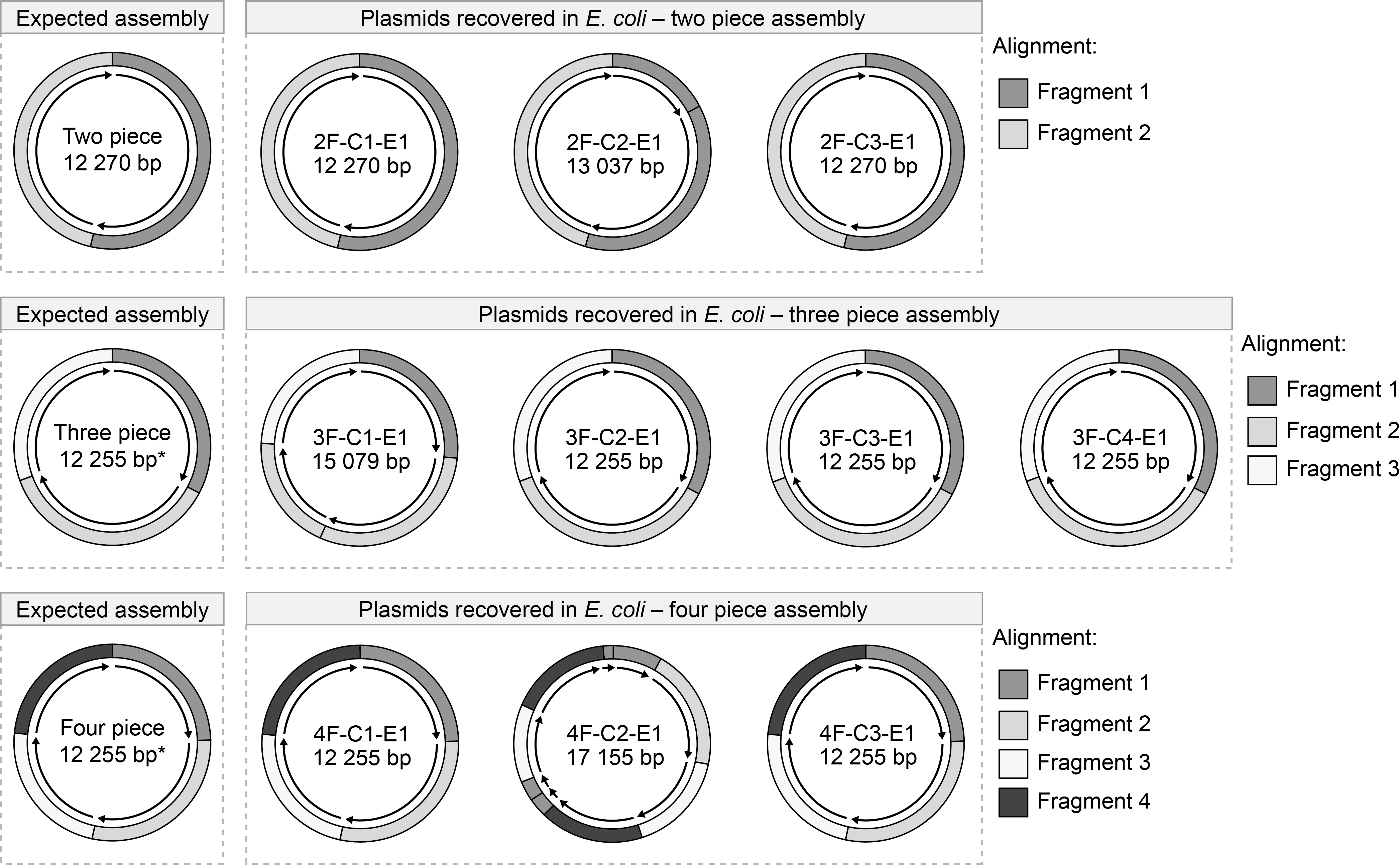

**Figure S8.** Sequencing of *P. tricornutum* HR-assembled episomes recovered from *E. coli* transformants. Fragment orientations are depicted by arrows and are relative to the DNA directionality in the original plasmid, pPtGE31_ShBle. * A 15-bp deletion was consistently observed across episomes recovered from the three- and four- piece assemblies. This was determined to be due to the accidental use of pPtGE31 as template DNA during amplification of these fragments. The expected episome is 15 bp smaller than that of the two-piece assembly because pPtGE31_ShBle contains a 15 bp insertion in a non-essential region of the episomal backbone.

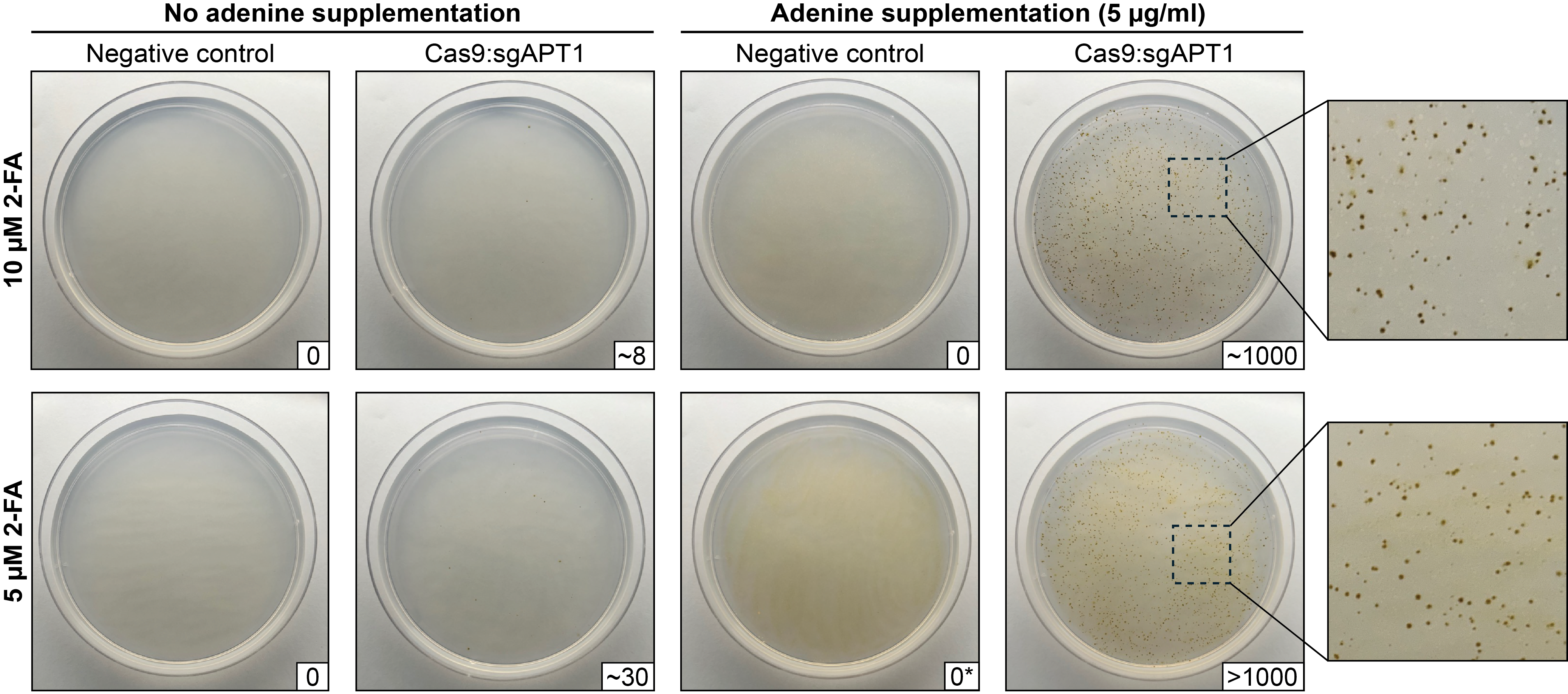

**Figure S9.** Electroporating an RNP complex (Cas9:sgAPT1) into spheroplasted *P. tricornutum* cells. Successful knockout of the PtAPT loci generates colonies that are resistant to 2-fluoroadenine (2-FA). One-fifth of the total reaction was plated across ½ salt L1 plates supplemented with 10 or 5 µM 2-FA and with or without an additional 5 µg/ml adenine. Plates were removed from the growth chamber and pictured 12 days post-electroporation to prevent breakthrough from ultraviolet irradiation degradation of the 2-FA. *No individual colonies were visible on this plate, however, a thin layer of algal growth is apparent.

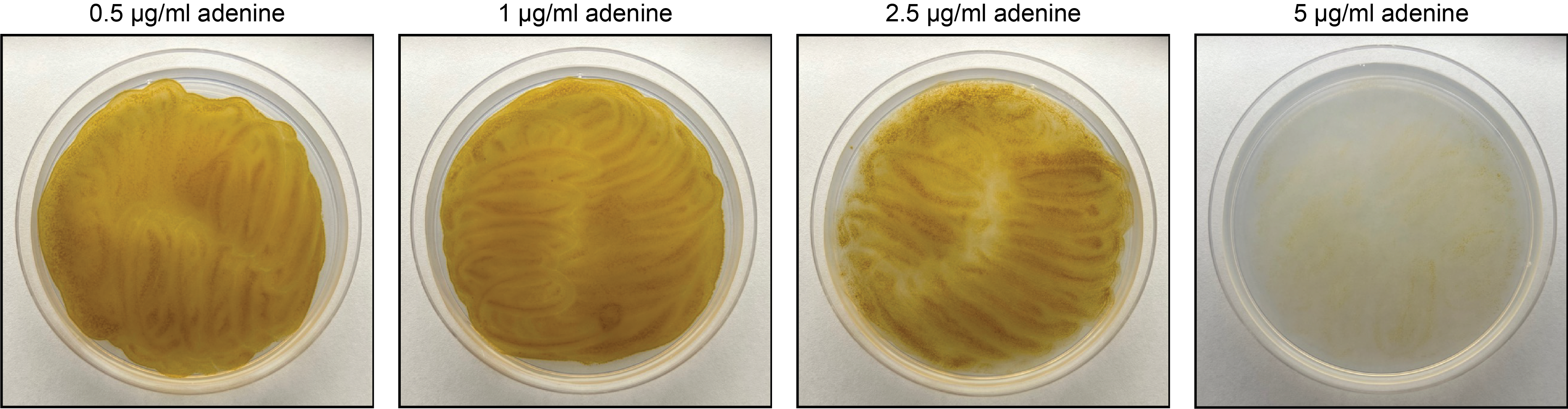

**Figure S10.** The growth of wild-type *P. tricornutum* on ½-salt L1 plates supplemented with differing amounts of adenine. Approximately 2.5 x 10^7^ cells (250 µl at a concentration of 1 x 10^8^ cells/ml) were spread onto each plate, which were then placed in a growth chamber for 1 week before being pictured.

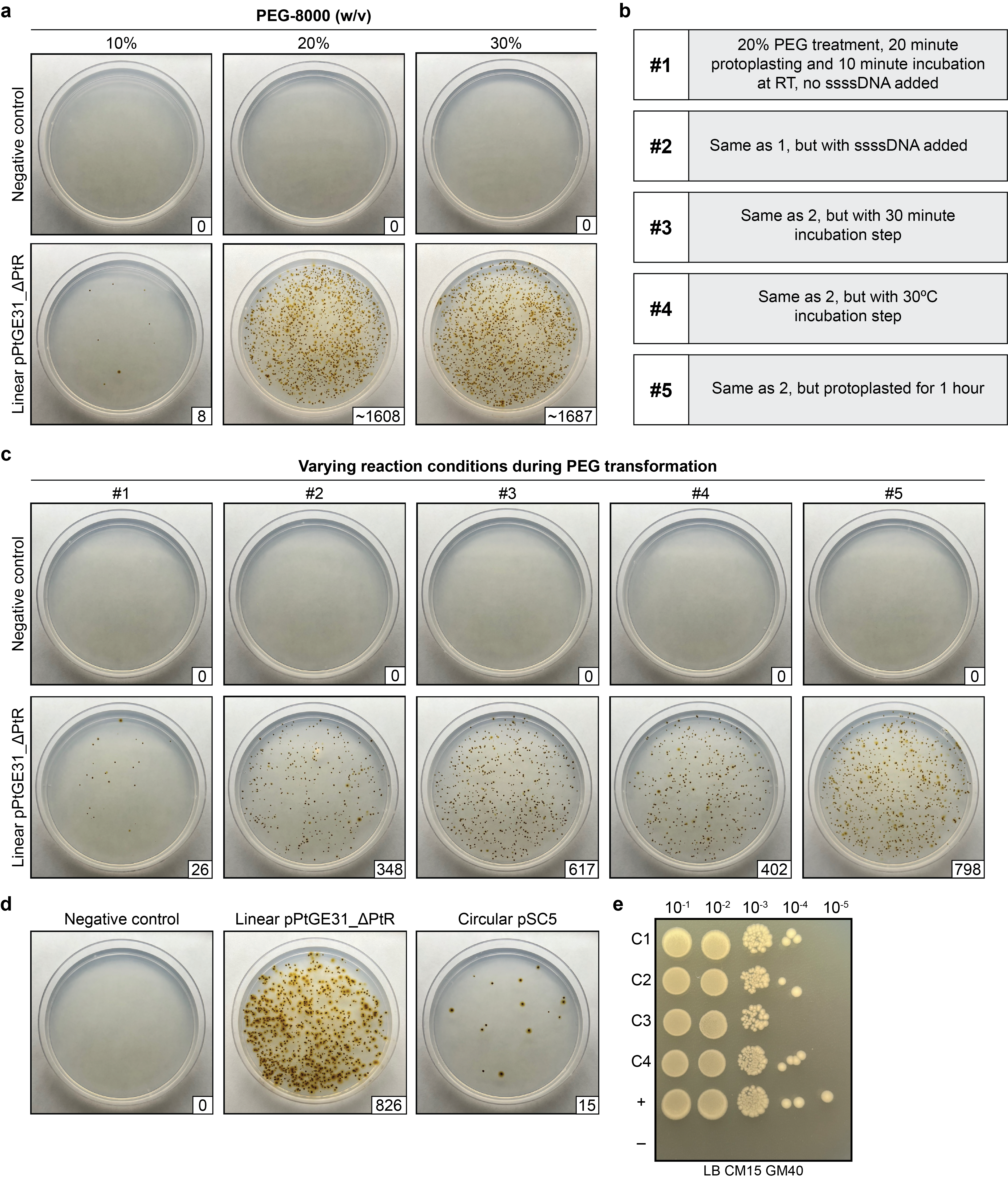

**Figure S11.** Establishing an efficient chemical transformation method for *P. tricornutum*. (**a**) Testing the transformation efficiency with different concentrations (w/v) of polyethylene glycol (PEG) 8000. (**b**) Five different experiments were carried out simultaneously, where one or more variables of the transformation method were changed. (**c**) The resulting transformation plates from the experimental conditions listed in b. (**d**) Transforming a 55.6 kb episome, pSC5, through the PEG method. (**e**) Testing the conjugation efficiency of *E. coli* strains harbouring episomes that had been recovered from *P. tricornutum* PEG transformants. Strains harbouring pSC5 or pSAP were used as the positive and negative controls, respectively. Dilutions of 10^-1^ to 10^-5^ were spot plated on LB plates supplemented with 15 µg/ml chloramphenicol and 40 µg/ml gentamicin. For experiments in panel a and d, a fourth of the PEG reaction was plated onto ¼-salt L1 plates supplemented with nourseothricin (100 µg/ml). In panel c, half of the PEG reaction was plated onto the same type of selection plates.

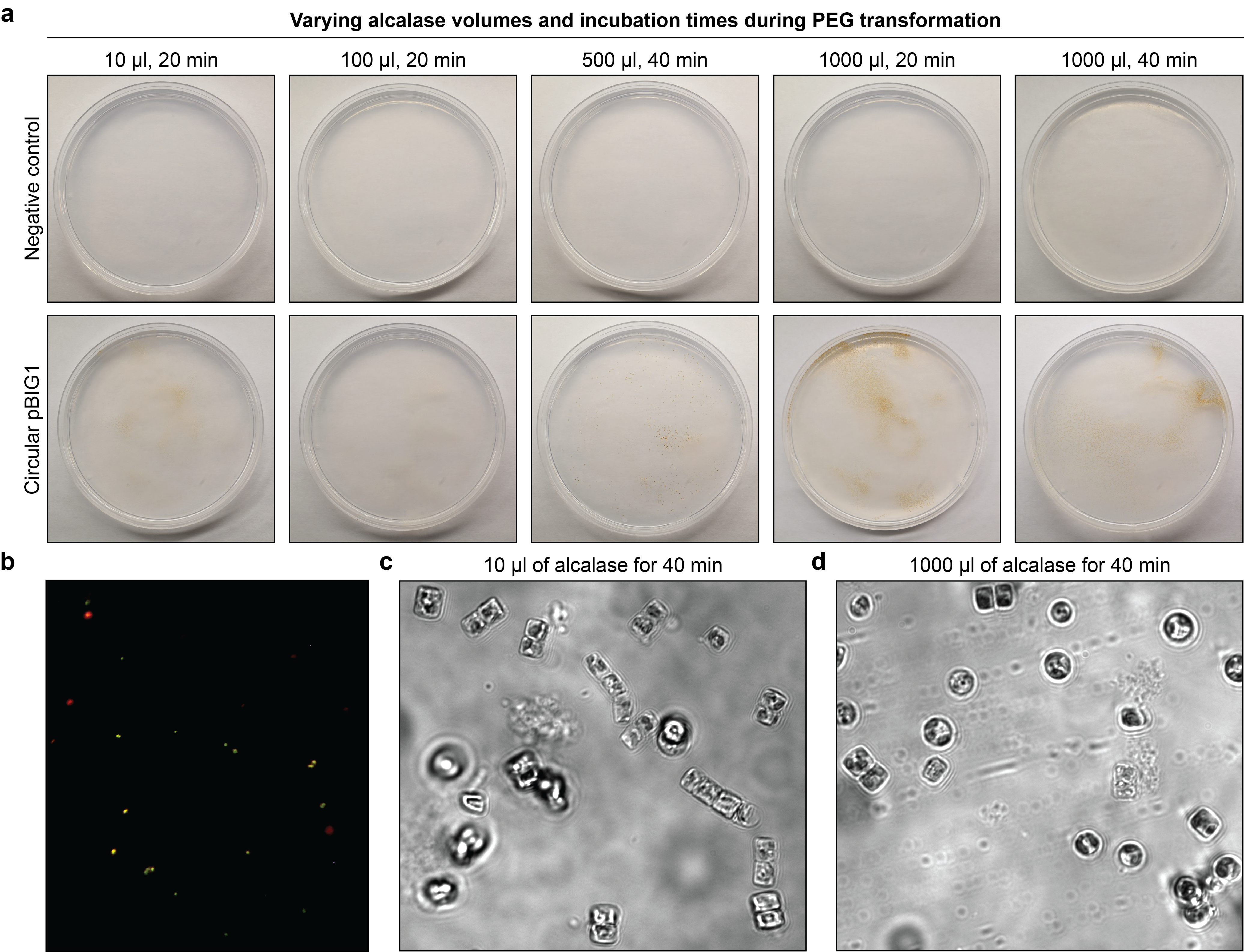

**Figure S12**. PEG transformation of *T. pseudonana*. (**a**) Harvested cells were subjected to differing amounts of alcalase and lengths of incubation during the protoplasting step of the PEG transformation method. Following recovery, half of the total reaction was plated on full-salt f/2 plates supplemented with 100 µg/ml nourseothricin. (**b**) Fluorescence microscopy screening of pooled PEG transformants. Red indicates autofluorescence and green indicates eGFP fluorescence. (**c**) Alcalase treatment of *T. pseudonana* with 10 µl or (**d**) 100 µl of alcalase for 40 minutes.

**Table S1.** Initial optimization of electroporation parameters for *P. tricornutum.* Each experiment used 1.0 – 2.0 x 10^6^ cells that had been harvested from liquid cultures. For experiments 60 to 74, transformants were plated on nourseothricin (NTC) and zeocin (ZEO) selection; NTC selection was used for all other experiments. The marker cassettes were generated through PCR amplification, whereas plasmids were purified from *E. coli*. Linear plasmid DNA was obtained through restriction digest with a single-cutter enzyme (exp. 1 – 6 and 18 – 22) or through PCR amplification (exp. 81 – 89). Intact circular plasmid DNA was used in experiments 77 – 79. **EE:** early exponential growth phase, **LS:** late stationary growth phase

| **Exp** | **DNA** | **Size**  (kb) | **Growth phase** | **Amount of DNA** (ng) | **Voltage**  (V) | **Capacitance** (µF) | **Resistance** (Ohms) | **CFUs** | **Total volume plated** |
| --- | --- | --- | --- | --- | --- | --- | --- | --- | --- |
| 1 | pPtGE27 | 8.6 | EE | 306 | 500 | 25 | 400 | 0 | 1/10 |
| 2 | pPtGE27 | 8.6 | EE | 306 | 500 | 25 | 400 | 0 | 1/10 |
| 3 | pPtGE27 | 8.6 | EE | 324 | 500 | 25 | 400 | 0 | 1/10 |
| 4 | pPtGE27 | 8.6 | EE | 390 | 500 | 25 | 400 | 0 | 1/10 |
| 5 | pPtGE27 | 8.6 | EE | 306 | 500 | 25 | 400 | 0 | 1/10 |
| 6 | pPtGE31 | 16.3 | EE | 468 | 500 | 25 | 400 | 0 | 1/10 |
| 7 | NAT cassette | 1.4 | EE | 606 | 500 | 25 | 400 | 0 | 1/10 |
| 8 | NAT cassette | 1.4 | EE | 792 | 500 | 25 | 400 | 0 | 1/10 |
| 9 | NAT cassette | 1.4 | EE | 462 | 500 | 25 | 400 | 0 | 1/10 |
| 10 | NAT cassette | 1.4 | EE | 462 | 500 | 50 | 400 | 2 | 1/10 |
| 11 | NAT cassette | 1.4 | EE | 462 | 500 | 50 | 150 | 0 | 1/10 |
| 12 | NAT cassette | 1.4 | EE | 462 | 500 | 25 | 400 | 0 | 1/10 |
| 13 | NAT cassette | 1.4 | EE | 462 | 500 | 50 | 400 | 2 | 1/10 |
| 14 | NAT cassette | 1.4 | EE | 462 | 500 | 50 | 150 | 0 | 1/10 |
| 15 | NAT cassette | 1.4 | EE | 462 | 500 | 25 | 400 | 0 | 1/10 |
| 16 | NAT cassette | 1.4 | EE | 462 | 500 | 50 | 400 | 6 | 1/10 |
| 17 | NAT cassette | 1.4 | EE | 462 | 500 | 50 | 150 | 0 | 1/10 |
| 18 | pPtGE27 | 8.6 | EE | 750 | 500 | 25 | 400 | 0 | 1/10 |
| 19 | pPtGE27 | 8.6 | EE | 750 | 500 | 50 | 400 | 2 | 1/10 |
| 20 | pPtGE27 | 8.6 | EE | 750 | 500 | 50 | 150 | 7 | 1/10 |
| 21 | pPtGE31 | 16.3 | EE | 660 | 500 | 25 | 400 | 0 | 1/10 |
| 22 | pPtGE31 | 16.3 | EE | 660 | 500 | 50 | 400 | 0 | 1/10 |
| 23 | NAT cassette | 1.4 | EE | 462 | 501 | 25 | 400 | 0 | 1/10 |
| 24 | NAT cassette | 1.4 | EE | 462 | 500 | 50 | 400 | 0 | 1/10 |
| 25 | NAT cassette | 1.4 | EE | 774 | 500 | 50 | 150 | 0 | 1/10 |
| 26 | NAT cassette | 1.4 | EE | 774 | 500 | 50 | 150 | 0 | 1/10 |
| 27 | NAT cassette | 1.4 | EE | 1320 | 500 | 50 | 400 | 1000+ | 1/10 |
| 28 | NAT cassette | 1.4 | EE | 1320 | 500 | 50 | 400 | 1000+ | 1/10 |
| 29 | NAT cassette | 1.4 | EE | 1320 | 500 | 50 | 150 | 657 | 1/10 |
| 30 | NAT cassette | 1.4 | EE | 1320 | 500 | 50 | 150 | 84 | 1/10 |
| 31 | NAT cassette | 1.4 | EE | 1320 | 486 | 75 | 150 | 84 | 1/10 |
| 32 | NAT cassette | 1.4 | EE | 1320 | 486 | 75 | 150 | 122 | 1/10 |
| 33 | NAT cassette | 1.4 | EE | 3300 | 500 | 50 | 150 | 389 | 1/10 |
| 34 | NAT cassette | 1.4 | EE | 3300 | 500 | 50 | 150 | 214 | 1/10 |
| 35 | NAT cassette | 1.4 | EE | 3300 | 487 | 75 | 150 | 67 | 1/10 |
| 36 | NAT cassette | 1.4 | EE | 840 | 500 | 50 | 400 | 0 | 1/10 |
| 37 | NAT cassette | 1.4 | EE | 840 | 500 | 50 | 400 | 0 | 1/10 |
| 38 | NAT cassette | 1.4 | EE | 840 | 500 | 50 | 150 | 0 | 1/10 |
| 39 | NAT cassette | 1.4 | EE | 840 | 500 | 50 | 150 | 0 | 1/10 |
| 40 | CAT/NAT cassette | 2.2 | EE | 2172 | 500 | 50 | 400 | 1 | 1/10 |
| 41 | CAT/NAT cassette | 2.2 | EE | 2172 | 500 | 50 | 400 | 49 | 1/10 |
| 42 | CAT/NAT cassette | 2.2 | EE | 2172 | 500 | 50 | 400 | 37 | 1/10 |
| 43 | CAT/NAT cassette | 2.2 | EE | 432 | 500 | 50 | 400 | 11 | 1/10 |
| 44 | CAT/NAT cassette | 2.2 | EE | 432 | 500 | 50 | 400 | 10 | 1/10 |
| 45 | CAT/NAT cassette | 2.2 | EE | 432 | 500 | 50 | 400 | 5 | 1/10 |
| 46 | ShBle/NAT cassette | 2.0 | EE | 912 | 500 | 50 | 400 | 0 | 1/10 |
| 47 | CAT/NAT cassette | 2.2 | EE | 2172 | 500 | 50 | 400 | 0 | 1/10 |
| 48 | NAT cassette | 1.4 | EE | 834 | 500 | 50 | 400 | 0 | 1/10 |
| 49 | ShBle/NAT cassette | 2.0 | EE | 312 | 500 | 50 | 400 | 0 | 1/10 |
| 50 | CAT/NAT cassette | 2.2 | EE | 270 | 500 | 50 | 400 | 0 | 1/10 |
| 51 | NAT cassette | 1.4 | EE | 180 | 500 | 50 | 400 | 0 | 1/10 |
| 52 | ShBle/NAT cassette | 2.0 | EE | 906 | 500 | 50 | 400 | 25 | 1/10 |
| 53 | ShBle/NAT cassette | 2.0 | EE | 906 | 500 | 50 | 400 | 28 | 1/10 |
| 54 | ShBle/NAT cassette | 2.0 | EE | 906 | 500 | 50 | 400 | 18 | 1/10 |
| 55 | ShBle/NAT cassette | 2.0 | EE | 240 | 500 | 50 | 400 | 5 | 1/10 |
| 56 | ShBle/NAT cassette | 2.0 | EE | 240 | 500 | 50 | 400 | 6 | 1/10 |
| 57 | NAT cassette | 1.4 | EE | 2622 | 500 | 50 | 400 | 2 | 1/10 |
| 58 | CAT/NAT cassette | 2.2 | EE | 588 | 500 | 50 | 400 | 38 | 1/10 |
| 59 | ShBle/NAT cassette | 2.0 | EE | 1278 | 500 | 50 | 400 | 120 | 1/10 |
| 60 | ShBle/NAT cassette | 2.0 | EE | 2718 | 500 | 50 | 400 | 790 (NTC)  47 (ZEO) | 1/10 |
| 61 | ShBle/NAT cassette | 2.0 | EE | 2718 | 500 | 50 | 400 | 333 (NTC)  20 (ZEO) | 1/10 |
| 62 | ShBle/NAT cassette | 2.0 | EE | 2718 | 500 | 50 | 400 | 306 (NTC)  18 (ZEO) | 1/10 |
| 63 | ShBle/NAT cassette | 2.0 | EE | 2400 | 500 | 50 | 400 | 1322 (NTC)  195 (ZEO) | 1/10 |
| 64 | ShBle/NAT cassette | 2.0 | EE | 2400 | 500 | 50 | 400 | 782 (NTC)  107 (ZEO) | 1/10 |
| 65 | ShBle/NAT cassette | 2.0 | EE | 2400 | 500 | 50 | 400 | 0 (NTC)  0 (ZEO) | 1/10 |
| 66 | ShBle/NAT cassette | 2.0 | EE | 1734 | 500 | 50 | 400 | 547 (NTC)  82 (ZEO) | 1/10 |
| 67 | ShBle/NAT cassette | 2.0 | EE | 1734 | 500 | 50 | 400 | 614 (NTC)  93 (ZEO) | 1/10 |
| 68 | ShBle/NAT cassette | 2.0 | EE | 1734 | 500 | 50 | 400 | 479 (NTC)  53 (ZEO) | 1/10 |
| 69 | ShBle/NAT cassette | 2.0 | EE | 960 | 500 | 50 | 400 | 218 (NTC)  73 (ZEO) | 1/10 |
| 70 | ShBle/NAT cassette | 2.0 | EE | 960 | 500 | 50 | 400 | 95 (NTC)  12 (ZEO) | 1/10 |
| 71 | ShBle/NAT cassette | 2.0 | EE | 960 | 500 | 50 | 400 | 56 (NTC)  14 (ZEO) | 1/10 |
| 72 | ShBle/NAT cassette | 2.0 | EE | 960 | 500 | 50 | 400 | 53 (NTC)  7 (ZEO) | 1/10 |
| 73 | ShBle/NAT cassette | 2.0 | EE | 960 | 500 | 50 | 400 | 52 (NTC)  6 (ZEO) | 1/10 |
| 74 | ShBle/NAT cassette | 2.0 | EE | 960 | 500 | 50 | 400 | 141 (NTC)  13 (ZEO) | 1/10 |
| 75 | NAT cassette | 1.6 | EE | 1080 | 500 | 50 | 400 | 214 | 1/10 |
| 76 | ShBle cassette | 1.5 | EE | 1080 | 500 | 50 | 400 | 16 | 1/10 |
| 77 | PtMt | 94.9 | EE | 849.6 | 500 | 50 | 400 | 0 | 1/10 |
| 78 | ptMt Min | 59.8 | EE | 2688 | 500 | 50 | 400 | 0 | 1/10 |
| 79 | pDMI2 | 16.4 | EE | 4056 | 500 | 50 | 400 | 0 | 1/10 |
| 80 | pPtGE31_ΔPtR | 10.9 | EE | 500 | 500 | 50 | 400 | 3 | 1/5 |
| 81 | pPtGE31_ΔPtR | 10.9 | EE | 500 | 500 | 50 | 400 | 5 | 1/5 |
| 82 | pPtGE31_ΔPtR | 10.9 | LS | 500 | 500 | 50 | 400 | 272 | 1/2 |
| 83 | pPtGE31_ΔPtR | 10.9 | LS | 500 | 500 | 50 | 400 | 265 | 1/2 |
| 84 | pPtGE31_ΔPtR | 10.9 | LS | 500 | 500 | 50 | 400 | 198 | 1/2 |
| 85 | pPtGE31_ΔPtR | 10.9 | LS | 500 | 500 | 50 | 400 | 35 | 1/2 |
| 86 | pPtGE31_ΔPtR | 10.9 | LS | 500 | 500 | 50 | 400 | 26 | 1/2 |
| 87 | pPtGE31_ΔPtR | 10.9 | LS | 500 | 500 | 50 | 400 | 37 | 1/2 |
| 88 | pPtGE31_ΔPtR | 10.9 | LS | 500 | 500 | 50 | 400 | 7 | 1/2 |
| 89 | pPtGE31_ΔPtR | 10.9 | LS | 500 | 500 | 50 | 400 | 256 | 1/2 |

**Table S2**. Transformation efficiencies for *P. tricornutum* cells cultured in liquid media or on agar plates, with or without alcalase treatment. Data were analyzed from five biological replicates, with there being an average of 1.8 x 10^8^ cells per transformation reaction. The values for mean CFUs were rounded to the nearest whole integer; all other values were rounded to three significant digits.

| **Media and treatment** | **Mean CFUs per reaction** | **Standard error of the mean** | **Efficiency** |
| --- | --- | --- | --- |
| Liquid, untreated | 76 | 19.9 | 4.22 x 10^7^ |
| Liquid, treated | 280 | 137.8 | 1.56 x 10^6^ |
| Plated, untreated | 108 | 43.0 | 6.02 x 10^7^ |
| Plated, treated | 21,495 | 5.44 x 10^3^ | 1.19 x 10^4^ |

**Table S3**. Transformation efficiencies for *P. tricornutum* cells transformed with differing amounts of PCR-amplified pPtGE31_ ΔPtR. Data were analyzed from four biological replicates, with there being an average of 1.24 x 10^8^ cells per transformation reaction. The values for mean CFUs per reaction and per ng of DNA were rounded to the nearest whole integer; the standard error of the mean values were rounded to three significant digits.

| **Amount of DNA** | **Mean CFUs per reaction** | **Standard error of the mean** | **CFUs per ng DNA** |
| --- | --- | --- | --- |
| 1 ng | 7 | 4.04 | 7 |
| 10 ng | 372 | 136 | 37 |
| 100 ng | 1545 | 536 | 16 |
| 500 ng | 13,109 | 5.38 x 10^3^ | 26 |
| 1000 ng | 16,629 | 6.68 x 10^3^ | 17 |

**Table S4**. Transformation efficiencies for *P. tricornutum* cells transformed with circular and linear pSC5. Data were analyzed from three biological replicates, with there being an average of 0.908 x 10^8^ cells per transformation reaction. The values for mean CFUs per reaction were rounded to the nearest whole integer; the standard error of the mean and efficiency values were rounded to three significant digits.

| **DNA** | **Mean CFUs per reaction** | **Standard error of the mean** | **Efficiency** |
| --- | --- | --- | --- |
| Circular pSC5 | 4 | 3.06 | 4.40 x 10^8^ |
| Linear pSC5 | 12 | 2.31 | 1.32 x 10^7^ |

**Table S5.** Passaging of *P. tricornutum* colonies transformed with two fragments, both individually and simultaneously. Transformants were initially replated on ¼-salt L1 plates supplemented with the same antibiotic selection as the original transformation plate (N = 100 µg/ml nourseothricin, Z = 100 µg/ml zeocin). On the second passage, colonies were repatched onto ¼-salt L1 plates containing the alternative antibiotic selection to screen for the presence of both resistance marker cassettes. Fragment 1 carries the zeocin-resistance marker, whereas fragment 2 carries the nourseothricin-resistance marker. Single fragment transformants were only plated on one type of selection plate.

| **Fragment(s)** | **Initial colonies** | | **First passage** | | **Second passage** | |
| --- | --- | --- | --- | --- | --- | --- |
|  | **N** | **Z** | **N 🡪 N** | **Z 🡪 Z** | **N 🡪 N 🡪 Z** | **Z 🡪 Z 🡪 N** |
| Fragment 1 | N/A | 12 | N/A | 10/12 | N/A | 0/10 |
| Fragment 2 | 81 | N/A | 12/20 | N/A | 0/12 | N/A |
| Both fragments | 250 | 16 | 100/100 | 16/16 | 79/100 | 15/16 |

**Table S6.** Passaging of *P. tricornutum* colonies co-transformed with pPtGE31_ΔPtR and Cas9:sgAPT1. One-fifth of the transformation volume was plated across two selection types: (1) ½-salt L1 plates supplemented with 5 µg/ml adenine and 10 µM 2-FA, and (2) ½-salt L1 plates supplemented with 1 µg/ml adenine and 100 µg/ml nourseothricin. Fifty colonies were first passaged from each selection plate onto the same selection type. On the second passage, colonies were repatched onto the other selection type.

| **Selection type** | **Initial colonies** | **First passage** | **Second passage** |
| --- | --- | --- | --- |
| NAT | 80 | 50/50 | 7/50 |
| 2-FA | 50 | 48/50 | 6/50 |

**Table S7**. Oligonucleotides used in this study.

| **Amplicon** | **Primers (5′ to 3′)** | **Template** | **Length (bp)** |
| --- | --- | --- | --- |
| **Amplification of fragments used in electroporation experiments** | | | |
| NAT marker | **F:**actagcttgattgggatatctcgctcgtgc  **R:**aaaccaaagcggagtgactgcaactaatg | pPtGE31 | 1431 |
| Linearized pPtGE31 plasmid | **F:**tcgagctggttgccctcgccgctgggctggcggccgtctatggccctgcaaacgcgccag  **R:**aaaccaaagcggagtgactgcaactaatgaaaattaaatttttaaggaaggatagagact  **Note**: we used phosphorylated and non-phosphorylated versions of these primers |  | 10,944 |
| **Primers used for screening transformation of *P. tricornutum* with linearized or circular pPtGE31** | | | |
| Termini region | **F:**actacctcgactttggctgg  **R:**atcatctgtgggaaactcgc | DNA isolated from algal colonies | 463 |
| Intact region of the backbone | **F:**cactgaagactgcgggattg  **R:**ggcttctcttatggcaaccg |  | 334 |
| **Primers used for two-piece NHEJ assembly of pPtGE31_ShBle in *P. tricornutum* (non-overlapping fragments)** | | | |
| F1 | **F:**attctcaaaatagcattttcgacatgcggtgctgatttca  **R:**gcaacagtttcaatggccagtcggagcatcaggtgtgg | pPtGE31_ShBle | 5483 |
| F2 | **F:**tcatgactatgatgtcatagttattgacagcgcgcctaac  **R:**cgccgatgctaatcattggcctctaaaaaaacggaagtatcaagaattccagacaaatcagg |  | 6787 |
| Screening junction A | **F:**ggactggactagcgtttggt  **F:**tcccgaatttgctcctccat  **R:**ctgttccaagtccgcaagtt | DNA isolated from algal and  *E. coli* colonies | 238 OR 359 |
| Screening junction B | **F:**gtgttttgctcgtggaaggt  **R:**tcccgaatttgctcctccat  **R:**ggactggactagcgtttggt |  | 553 OR 432 |
| **Primers used for two-piece HR assembly of pPtGE31_ShBle in *P. tricornutum* (overlapping fragments)** | | | |
| F1 | **F:** GGACTCCCGGACGTTCGT  **R:** TCCCGAATTTGCTCCTCCAT | pPtGE31_ShBle | 6626 |
| F2 | **F:** TTTGACTACACCTCCGCACT  **R:** GTCGCGAGCCCCATCAAC |  | 5964 |
| Junction between F1 and F2 | **F:** ATGTGCTGATTGTTCCCACG  **R:** AAGAGCGTTGATCAATGGCC | DNA isolated from algal and  *E. coli* colonies | 300 |
| **Primers used for three-piece HR assembly of pPtGE31_ShBle in *P. tricornutum* (overlapping fragments)** | | | |
| F1 | **F:** GGACTCCCGGACGTTCGT  **R:** CGCAGCCGTGTAACCGAG | pPtGE31_ShBle | 4106 |
| F2 | **F:** CTATCCGCGTGTGTACCTCT  **R:** GACACTGAATACGGGGCAAC | *pPtGE31 | 4669 |
| F3 | **F:** CTGTGGATTGCTGCTGTGTC  **R:** GTCGCGAGCCCCATCAAC | *pPtGE31 | 3971 |
| Junction between F1 and F2 | **F:** GGGGTTAGTTCGTCATCATTGA  **R:** GAGAGCAGGAAGAGCAAGGT | DNA isolated from algal and  *E. coli* colonies | 379 |
| Junction between F2 and F3 | **F:** CTCGGTGTGCGGTTGTATG  **R:** GGCCATCAAACCACGTCAAA |  | 298 |
| **Primers used for four-piece HR assembly of pPtGE31_ShBle in *P. tricornutum* (overlapping fragments)** | | | |
| F1 | **F:** GGACTCCCGGACGTTCGT  **R:** AGCAACATGGATCTCGCAGA | pPtGE31_ShBle | 3137 |
| F2 | **F:** CGCGAATCGTCCAGTCAAAC  **R:** TCCCGAATTTGCTCCTCCAT | *pPtGE31 | 3638 |
| F3 | **F:** TTTGACTACACCTCCGCACT  **R:** TGACCATGATTACGCCAAGC |  | 3030 |
| F4 | **F:** GTTGTAAAACGACGGCCAGT  **R:** GTCGCGAGCCCCATCAAC |  | 3084 |
| Junction between F1 and F2 | **F:** GGATCTGTCATGGCGGAAAC  **R:** ACTCTTCAATGCCTGCCGTA | DNA isolated from algal and  *E. coli* colonies | 273 |
| Junction between F2 and F3 | **F:** CGACTGGCCATTGAAACTGT  **R:** AAGAGCGTTGATCAATGGCC |  | 400 |
| Junction between F3 and F4 | **F:** CTCTTCGCTATTACGCCAGC  **R:** TCACTCATTAGGCACCCCAG |  | 333 |
| **Primers used for screening allele 1 and 2 at the PtAPT loci, obtained from the supplementary data in Serif et al. (2018)** | | | |
| Allele non-specific primers | **F:** TCCTCTGTTCGACTGCTCGT  **R:** ACCAAACTAACGGCCGACC | DNA isolated from algal colonies | ~750 |
| Allele 1 primers | **F:** CGACATACCCGGAGGTTAGTT  **R:** CGCATCATTATAAAGGGTTTATTCAA |  | ~611 |
| Allele 2 primers | **F:** CGACATACCCGGAGGTTAGTG  **R:** CGCATCATTATAAAGGGTTTATTCAG |  | ~611 |
| **Amplification of fragments to generate pPtGE31_ShBle via yeast assembly**  Capitalized base pairs share homology to pPtGE31  Lower-case base pairs share homology to p0521S | | | |
| ShBle ORF with overhangs to insert into pPtGE31 | **F:**AATTTAATTTTCATTAGTTGCAGTCACTCCGCTTTGGTTTcaggattagtgcaattcgagttgaatcactgggaaaaaca  **R:**TAGACGGCCGCCAGCCCAGCGGCGAGGGCAACCAGCTCGAaaaccaaagcggagtgactgcaactaatgaaaattaaatt | p0521S | 1406 |
| First half of pPtGE31, has an overhang for the ShBle ORF | **F:**aatttaattttcattagttgcagtcactccgctttggtttTCGAGCTGGTTGCCCTCGCCGCTGGGCTGGCGGCCGTCTA  **R:**GTAAAGACAGATAAACGTAGACTAAAACGAGGTCGCATCAGGGTGCTGGCTTTTCAAGTT | PtGE31_ΔPtR | 5942 |
| Second half of pPtGE31, has an overhang for the ShBle ORF | **F:**CGACATCAGTTTGCTCCTGGAGCGACAGTATTGTATAAGGGCGATAAAATGGTGCTTAAC  **R:**tgtttttcccagtgattcaactcgaattgcactaatcctgAAACCAAAGCGGAGTGACTGCAACTAATGAAAATTAAATT |  | 5260 |

*The template pPtGE31 was accidentally used when amplifying fragments F2 and F3 of the three-piece assembly and fragments F2, F3, and F4 of the four-piece assembly. This made no functional changes to the assembled episome.
